# GluN2B-mediated regulation of silent synapses for receptor specification and addiction memory

**DOI:** 10.1101/2024.05.19.594887

**Authors:** Hyun Jin Kim, Sangjun Lee, Gyu Hyun Kim, Kibong Sung, Taesik Yoo, Jung Hyun Pyo, Hee-Jung Jo, Sanghyeon Lee, Hyun-Young Lee, Jung Hoon Jung, Kea Joo Lee, Joung-Hun Kim

**Affiliations:** Department of Life Sciences, Pohang University of Science and Technology (POSTECH), Pohang, Gyungbuk, Republic of Korea; Salk Institute for Biological Studies, La Jolla, CA, USA; Neural Circuits Research Group, Korea Brain Research Institute (KBRI), Daegu, Republic of Korea; Department of Biomedical Sciences, Seoul National University College of Medicine, Seoul, Republic of Korea; Program in Neurosciences & Mental Health, Hospital for Sick Children, Toronto, ON, Canada

## Abstract

Psycho-stimulants including cocaine elicit stereotyped, addictive behaviors. Re-emergence of silent synapses containing only NMDA-type glutamate receptors (NMDARs) is a critical mediator of addiction memory and seeking behaviors. Despite the predominant abundance of GluN2B-containing NMDARs in silent synapses, their operational mechanisms are not fully understood. Using conditional depletion/deletion of GluN2B at D1-expressing accumbal medium spiny neurons, we examine the synaptic and behavioral actions that silent synapses incur after repeated exposure to cocaine. GluN2B ablation reduces the proportion of silent synapses, but some of them can persist by substitution to GluN2C, which drives the aberrantly-facilitated synaptic incorporation of calcium-impermeable AMPA-type glutamate receptors (AMPARs). The resultant precocious maturation of silent synapses impairs addiction memory but elevates locomotor activity, which can be normalized by blockade of calcium-impermeable AMPAR trafficking. Collectively, GluN2B supports the competence of cocaine-induced silent synapses for specifying subunit composition of AMPARs, and thereby expression of addition memory and the related behaviors.

## Introduction

Neural circuits and the physiological mechanisms underlying addiction memory and the associated behaviors to psycho-stimulants such as cocaine, have been extensively explored(Everitt & Robbins, 2005). For instance, the mesocorticolimbic system is regulated by dopamine that can elaborate and modulate synaptic plasticity, which shapes the acquisition and maintenance of addiction memory(Brown *et al*, 2011; Crittenden & Graybiel, 2011; Nestler, 2013; Pascoli *et al*, 2015). A significant portion of dopaminergic projections originating from the ventral tegmental area extends to the striatum where they innervate the medium spiny neurons (MSNs) expressing either dopamine receptor D1 (Drd1) or D2 (Drd2)(Gerfen & Surmeier, 2011). It has been believed that the dopamine-dependent regulation of synapses in MSNs of the nucleus accumbens (NAc) can adjust or determine development and intensification of addiction memory and seeking behaviors toward cocaine(Russo *et al*, 2010).

Neural innervations from the basolateral amygdala (BLA) to the NAc undergo synaptic plasticity leading to motivated behaviors toward cocaine as well as cue-induced reinstatement: selective enhancement of transmission strength from the BLA to Drd1-expressing MSNs (D1-MSNs) after chronic cocaine exposure(Kwon *et al*, 2015; MacAskill *et al*, 2014). Consistently, stimulation of the BLA-NAc pathway alone amplified the magnitude of addiction-related behaviors including locomotor sensitization (MacAskill *et al*., 2014; Namburi *et al*, 2015) and conditioned place preference (CPP)(de Guevara-Miranda *et al*, 2016), while activation of cortical projections did not do so(Stuber *et al*, 2011). Furthermore, cell type-specific output activity of NAc D1-MSNs exerted bidirectional effects onto extinction and reinstatement of cocaine seeking, along with CPP being disrupted by the modulation of actin polymerization within the NAc shell (NAcSh)(Gibson *et al*, 2018; Li *et al*, 2015). Hence, various forms of plasticity occurring at each type of MSN seemed to differentially modulate behavioral changes triggered by cocaine administration.

Generation and maturation of silent synapses lacking α-amino-3-hydroxy-5-methyl-4-isoxazolepropionic acid glutamate receptors (AMPARs)(Huang *et al*, 2009) in the NAc could be a major mediator for the development and intensification of cocaine-induced behaviors(Dong & Nestler, 2014; Huang *et al*., 2009; Lee & Dong, 2011). While silent synapses containing only N-methyl D-aspartate glutamate receptors (NMDARs) typically occurred at the early stages of neuronal development but diminish as neurons become mature, they reappeared after exposure to addictive substances including cocaine or amphetamine, which might represent a “rejuvenation” of neurons(Dong & Nestler, 2014).

Cocaine-generated silent synapses were preferentially comprised of GluN2B-containing NMDARs (GluN2B-NMDARs) and underwent maturation with incorporation of AMPARs (Wang *et al*, 2021), which can support expression of addiction memory and seeking behaviors(Huang *et al*., 2009; Wright *et al*, 2020). Despite their mechanistic and clinical importance, whether or not only GluN2B-NMDARs are able to constitute and maintain silent synapses, how they modulate synaptic modification, and which physiological significance they hold in the context of drug addiction remained to be clarified particularly at the circuit levels. Using a combination of region- and cell-type-specific depletion/deletion of GluN2B, electrophysiological recordings, electron and confocal microscopic analyses, and behavioral tests in wild-type and knock-in mice, we characterized the operational roles played by GluN2B-NMDARs for cocaine-induced silent synapses in NAc D1-MSNs as well as the physiological and behavioral roles after repeated administration of cocaine.

## Results

### Requirement of GluN2B for cocaine-induced silent synapses and addiction memory

It has been widely documented that physiological and structural changes of striatal circuits normally occur during extended abstinence periods (weeks to months) after chronic administration of addictive drugs(Wright *et al*., 2020). However, early forms of drug-evoked synaptic plasticity, if any, can be metaplasticity permissive for the late onset of adaptive physiological and behavioral changes with repetitive drug exposure. Thus, we monitored synaptic transmission and structural changes as early as several hours (< 3 hours) after completion of 5-day administration of cocaine.

AMPAR-lacking silent synapses appeared in D1-MSNs of the NAc after repeated administration of cocaine(Huang *et al*., 2009). Indeed, increased surface insertion of GluN2B-NMDARs resulted in the elevation of numbers of silent synapses(Brown *et al*., 2011; Huang *et al*., 2009; Lee & Dong, 2011)whereas the acute depletion prevented *de novo* formation of silent synapses(Huang *et al*., 2009). To explore the operational roles of silent synapses, we depleted GluN2B, specifically in D1-MSNs of the NAcSh. This was achieved through viral Cre-dependent expression of small hairpin RNA targeting GluN2B (shGluN2B, Figs. 1A and EV1A) (Kwon *et al*., 2015), the efficacy of which was validated via western blotting (Fig. EV1B). Importantly, AAV containing shGluN2B gave rise to an apparent reduction of GluN2B expression without any discernible effects on GluN2A or GluN1 expression (Figs. EV 1B-D). Moreover, the ablation of GluN2B was confirmed through quantification of GluN2B-positive puncta in D1-MSNs (Figs. 1B, and EV1E) and immuno-transmission electron microscope (EM)-based ultra-structural analysis at the single-synapse level (Figs. 1C, EV1F).

**Figure 1.**
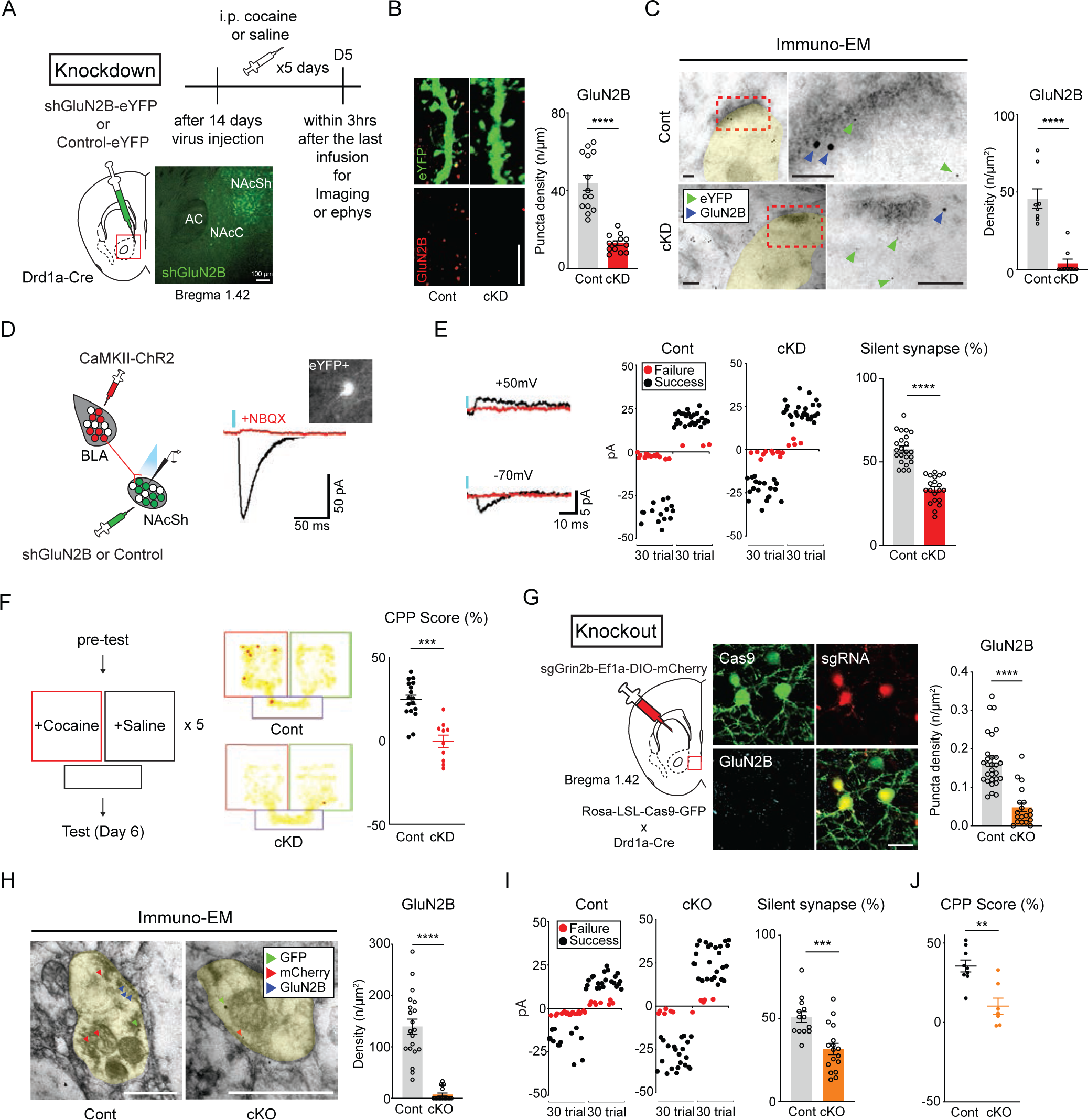
GluN2B are necessary for cocaine-induced silent synapses and addiction memory. **(A)** Schematic for shGluN2B or control virus injection in the NAcSh (left). A timeline for general experiments-cocaine or saline injections were made once every day (5 days, top, right) and expression of shGluN2B virus in NAcSh area (bottom, right). Scale bar = 100 μm. **(B)** Representative images of GluN2B-positive puncta (red) in D1-MSNs dendrites (green, left). GluN2B puncta density after cocaine administration is compared between mice that received either shGluN2B or control virus (n = 13 cells for cKD; n = 14 cells for control, right). Scale bar = 5 μm. **(C)** Immuno-electron microscopic images showing subcellular localization of GluN2B (18 nm gold particles, blue arrows) and eYFP for labeling D1-MSNs (6 nm gold particles, green arrows, left). GluN2B particle density is compared between cocaine/cKD and cocaine/control D1-MSNs (n = 9 cells for cKD; n = 8 cells for control, right). Scale bars = 100 nm. (**D)** Schematic for optogenetic stimulation of the BLA-NAcSh pathway (left). Representative EPSC traces from optical stimulation (blue bar) of BLA glutamatergic inputs with a picture with (red) or without NBQX (black) showing cKD D1-MSNs (an inserted image, right). (**E)** Sample traces of success (black trace) and failure (red trace) responses by the optical minimal stimulation for AMPAR-EPSCs (−70mV) and NMDAR-EPSCs (+50mV, left). Example plots of optical responses with minimal optical stimulation in designated groups (failure trials in red dots, successful trials in black dots, middle). The proportion of silent synapses is compared between cocaine-treated mice that previously received either shGluN2B or control virus (n = 20 cells for cKD; n = 23 cells for control group, right). **(F)** Schematic for conditional place preference (left). Heatmap traces of representative animals (middle). Quantified CPP scores to the cocaine chamber are compared between mice that previously received either shGluN2B or control virus (n = 10 mice for cKD; n = 17 mice for control group, right). **(G)** Schematic for cKO of GluN2B using CRISPR-Cas9 (left). Representative images for GluN2B (cyan) expression in D1-MSNs expressing Cas9-eGFP (green) and sgGrin2b-mCherry (red, middle). GluN2B puncta density after cocaine administration is compared between mice that previously received either sgGrin2b or control virus (n = 19 cells for cKO; n = 29 cells for control group, right). Scale bar = 20 μm. **(H)** Immuno-electron microscopic images showing subcellular localization of GluN2B (12 nm gold particles, blue arrows), mCherry (6 nm gold particles, red arrows) and eYFP for labeling D1-MSNs (18 nm gold particles, green arrows, left). GluN2B particle density is compared in the dendrites-labelled by eYFP between cocaine/cKO and cocaine/control group (n = 20 cells for cKO; n = 18 cells for control group, right). Scale bar = 100 nm. **(I)** Example plots of AMPAR- and NMDAR-EPSCs with minimal optical stimulation in designated groups (failure trials in red dots; successful trials in black dots, left). The proportion of silent synapses is compared between cocaine-treated Cas9-eGFP mice that previously received either sgGrin2b or control virus (n = 17 cells for cKO; n = 13 cells for the control group, right). (**J)** Quantified CPP scores in the cocaine chamber are compared between GluN2B cKO and control groups (n = 17 mice for cKO; n = 13 mice for the control group). Data are represented as mean ± SEM (error bars); **p < 0.01, ***p < 0.001, ****p < 0.0001 by two-tailed unpaired t test or Mann-Whitney test.

GluN2B depletion impaired cocaine-induced synaptic plasticity and seeking behaviors(Isaac *et al*, 1995). We also examined the physiological and behavioral impacts of GluN2B depletion particularly in the BLA-D1-MSNs pathway after 5-day cocaine exposure. To gain optogenetic control of the BLA-NAc pathway, we conducted optical stimulation of the BLA with an infusion of AAV encoding CamKIIα-ChR2-mCherry into the BLA of D1a-Cre mice that previously received shGluN2B virus (Fig. EV1G). Optical stimulation of BLA fibers projecting to the NAc was sufficient to evoke AMPAR-mediated excitatory postsynaptic currents (EPSCs) from eYFP-labelled D1-MNS with a delay time of 19.63 ± 4.65 ms, which were abolished by 2,3-dihydroxy-6-nitro-7-sulfamoyl-benzo[f]quinoxaline (NBQX, Fig. 1D). To obtain the proportion of silent synapses in D1-MSNs of mice that repeatedly received cocaine, we used a conventional method with minimal stimulation(Isaac *et al*., 1995; Stevens & Wang, 1995). Photo-stimulation intensity was adjusted to the point where EPSC failures and successes could be visually distinguished at −70 and +50 mV holding potentials as previously described(Graziane *et al*, 2016). The calculated percentages of cocaine-induced silent synapses were lower in conditional knock-down (cKD) mice than those in control group (Fig. 1E) while the same GluN2B depletion did not affect silent synapses in saline-treated mice (Fig. EV1H).

We used CPP to examine whether shGluN2B-mediated reduction of silent synapses could affect addiction memory(Tzschentke, 1998). Subject animals initially had no specific preference to any chambers before cocaine injection (Fig. EV1I), but they became to prefer the chamber where cocaine was injected. However, D1a-Cre mice that bilaterally received shGluN2B virus had no preference toward any chambers even after cocaine injection (Fig. 1F), indicating that shGluN2B-mediated reduction of silent synapses interfered with the formation of addiction memory.

To exclude the possibility that any observed effects were due to off-the-target effects of the used shRNA sequence, we also conditionally knocked-out (cKO) GluN2B in D1-MSNs using CRISPR-Cas9 system, the efficacy of which was validated via western blotting (Fig. EV2A). When AAV containing *sgGrin2b* was delivered to NAc areas of LSL-Cas9-GFP mice that had crossed with D1a-Cre mice, we detected co-localization of Cas9 and sgRNA (Fig. 1G). The puncta and particle density of GluN2B was lower in infected D1-MSNs than in control GFP-neurons (Figs. 1G, H and EV2B). Importantly, GluN2B cKO mice exhibited alteration of the proportion of silent synapses and cocaine-associated behaviors, which were almost identical to those of GluN2B cKD mice: a decrease in the proportion of silent synapses (Fig. 1I), impairment of CPP but normal appetitive memory to sucrose (Figs. 1J and EV2C). These data pointed to the essential roles of GluN2B both in generation or maintenance of silent synapses of D1-MSNs and in expression of addiction memory.

### Cocaine-induced structural and physiological changes facilitated by GluN2B ablation

In most cases, functional modification and maturation of synapses were associated with the structural changes in striatal neurons(Borczyk *et al*, 2019; Russo *et al*., 2010). We resorted on electron microscopies to thoroughly investigate structural features of D1-MSNs. Scanning electron microscopy (SEM) was utilized to examine subtle anatomical differences in dendritic structures between GluN2B-depleted and control D1-MSNs. We conducted 3-dimensional reconstruction of images obtained from SEM tomography, which achieved complex wiring diagrams of D1-MSN dendrites (Fig. 2A). Subsequent examination of reconstructed dendritic structures revealed morphological characteristics of dendritic spines of D1-MSNs accordingly to individual groups, cocaine-treated cKD (cocaine/cKD), cocaine-treated eYFP-expressing control (cocaine/control), saline-treated cKD (saline/cKD) and saline-treated control (saline/control) groups. We categorized the dendritic protrusions into four classes based on their shape and appearance(Dickstein *et al*, 2016; Rodriguez *et al*, 2008; Yuste, 2023): filopodia, thin spines, mushroom spines, and stubby spines (detailed in Methods). Our subsequent analysis indicated that a significant increase in the proportion of mushroom and stubby spines, indicative of mature synapses, was manifest in cocaine/cKD compared with cocaine/control group while much less filopodia and thin spines, normally considered to be immature synapses, were seen in cocaine/cKD (Fig. 2B). However, we failed to detect differences in general spine density across groups (Fig. EV3A) and in mature structures between groups without cocaine treatment (Fig. EV3B) in the time point for our observation. We also measured dimensions of each type of protrusions. The width of protrusion heads and the areas of postsynaptic density (PSD) were larger in cocaine/cKD group compared with those in cocaine/control group. Interestingly, we could frequently detect the increased numbers of branched spines in cocaine/cKD group (Fig. 2C), whereas the volume of spines and the numbers of synaptic vesicles (SVs) per unit area of the PSD and perforated PSD did not differ between groups (Figs. EV3C and D). Thus, SEM-based anatomical analysis revealed that the ablation of GluN2B elevated the number of morphologically-defined mature synapses while it did not affect the general spine density yet in several hours after repeated cocaine infusion.

**Figure 2.**
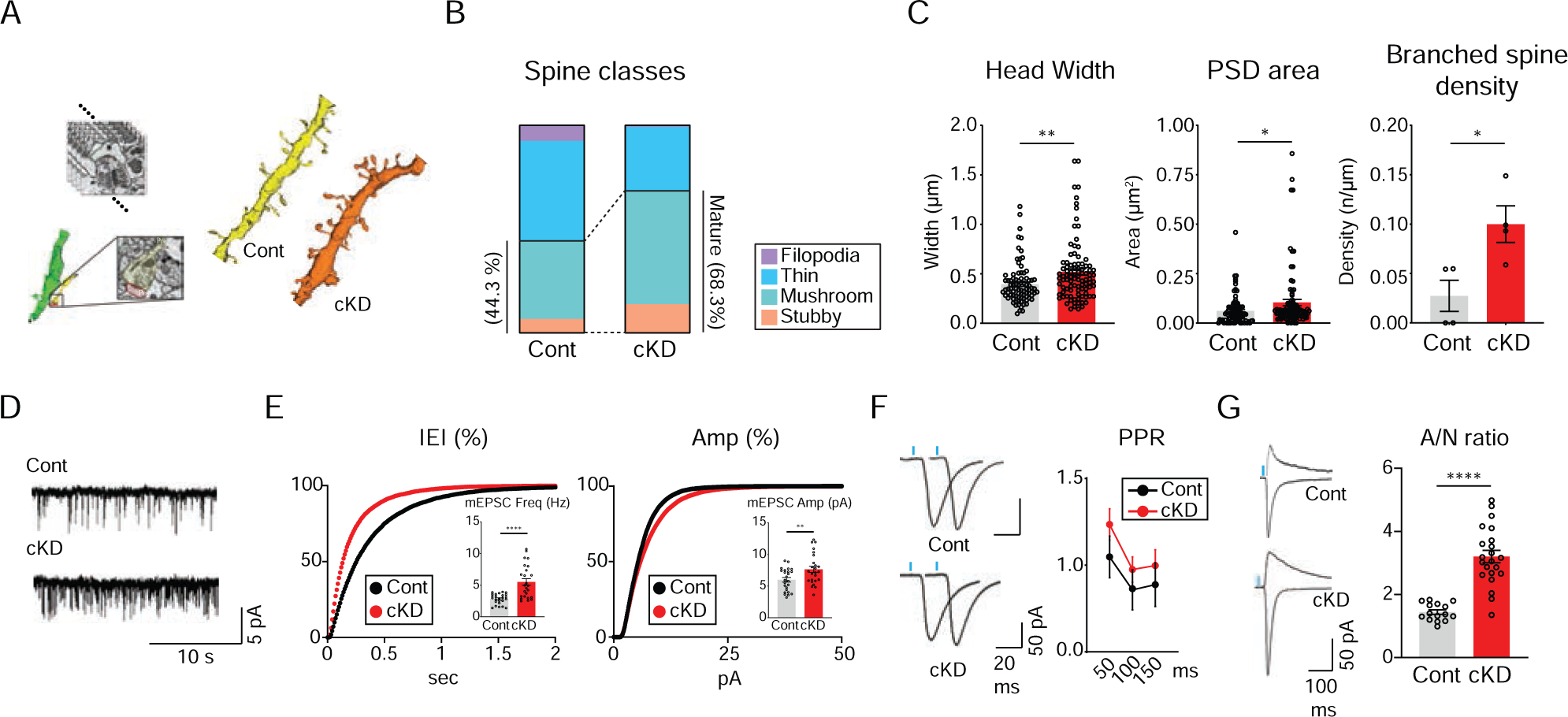
Structural and functional maturation of synapses are accelerated by GluN2B depletion. **(A)** A chematic for the 3-dimensional reconstruction from serial SEM images (left). Representative illustrations of reconstructed dendrites from cocaine-treated cKD and control D1-MSNs (right). **(B)** The proportion of morphologically-categorized spine classes between cocaine-treated mice that previously received either shGluN2B (cocaine/cKD, n = 79 spines from 3 mice) or control virus (cocaine/control, n = 85 spines mice from 2 mice). (**C)** Analysis of averaged spine head widths, PSD areas, and density of branched spines are compared between cocaine/cKD and cocaine/control groups. (head widths and PSD areas, n = 85 units for cocaine/control, n = 95 units for cocaine/cKD; density of branched spines, n = 4 units for cocaine/control, n = 4 units or cocaine/cKD). (**D)** Sample traces of mEPSCs from D1-MSNs of cocaine-treated mice that previously received either shGluN2B or control virus. (**E)** A cumulative probability plot of mEPSC inter-event intervals (IEIs) and an inserted summary histogram of quantified mEPSC frequency after cocaine exposure (n = 25 cells each for control and cKD groups, left). A cumulative probability plot of mEPSC amplitudes and a summary histogram for quantified mEPSC amplitudes after cocaine exposure (n = 25 cells each for cocaine/control and cocaine/cKD groups, right). (**F)** Sample traces of optically-evoked EPSCs with stimulation of 50-ms interval in D1-MSNs of saline-treated mice that previously received either shGluN2B or control virus (left). Measurement of paired-pulse ratios (PPRs) obtained with stimulation of 50-, 100-, and 150-ms intervals in D1-MSNs from cocaine-treated mice that previously received either shGluN2B or control virus (n = 9 cells for cKD; n = 11 cells for control groups, right). **(G)** Sample traces of AMPAR-(at −70 mV) and NMDAR-(+50 mV) EPSCs in cocaine-treated D1-MSNs (left). A/N ratios are compared between cocaine-treated mice that previously received either shGluN2B or control virus (n = 22 cells for cKD; n = 15 cells for control groups, right). Data are represented as mean ± SEM (error bars); *p < 0.05, **p < 0.01, ****p < 0.0001 by two-tailed unpaired t test or Mann-Whitney test.

We monitored miniature excitatory postsynaptic currents (mEPSCs) to assess basal transmission in D1-MSNs. Consistent with our anatomical data, we detected significant increases in mEPSC frequency and amplitude in cocaine/cKD group compared with those in cocaine/control group (Figs. 2D, E and EV3E-G). However, paired-pulse ratios (PPRs) of evoked EPSCs did not differ among groups (Figs. 2F and EV3H). On the other hand, ratios of AMPAR/NMDAR-EPSCs (A/N ratios) significantly increased in cocaine/cKD group compared with those in cocaine/control group (Figs. 2G and EV3I). These results supported the notion that D1-MSNs lacking GluN2B would have postsynaptic changes with increases in AMPAR-mediated components. Combined with anatomical results, these physiological data suggested that the observed decrease in the proportion of silent synapses would be due to facilitated maturation/un-silencing of silent synapses rather than impairment in the generation of silent synapses *per se*, assuming that silent synapses are precursors of AMPAR-containing functional synapses.

### GluN2B ablation-mediated recruitment of calcium-impermeable GluA2-AMPARs

It was previously shown that GluN2B deletion increased the surface expression of AMPARs in human embryonic kidney cells or hippocampal neurons *in vitro*(Kim *et al*, 2005). If this also occurred in NAc D1-MSNs, decreased proportion of silent synapses and increased occurrence of functional types of synapses that we have observed could be attributed to premature un-silencing of silent synapses due to promoted incorporation of AMPARs. It has been widely believed that recruitment of calcium-permeable (CP) AMPARs could mediate maturation of silent synapses in D1-MSNs after cocaine exposure and underlie incubation of addiction memory and craving traits(Brown *et al*., 2011; Lee & Dong, 2011; Lee *et al*, 2013). However, we detected no rectification of AMPAR-EPSCs that CP-AMPARs normally displayed at depolarized holding potentials(Henley & Wilkinson, 2016) (Figs. EV4A and B). These results would be seemingly consistent with the previous reports showing that the rectification of AMPAR-EPSCs appeared during extended abstinence period (> 7 days) after repeated cocaine infusion(Boudreau & Wolf, 2005; Lee *et al*., 2013). Given our anatomical and physiological data suggesting precocious maturation of D1-MSNs in cocaine/cKO mice, however, the absence of AMPAR-EPSC rectification could be due to facilitated recruitment of other types of AMPARs rather than CP-AMPARs. To this end, we took advantage of non-stationary fluctuation analysis (NSFA) from optically-evoked AMPAR-EPSCs(Sun *et al*, 2007) to approximate single-channel currents of individual AMPARs of D1-MSNs. Our NSFA revealed that the single-channel currents in cocaine/cKD group were significantly smaller than those in cocaine/control group (Figs. 3A and 4C), arguing against the incorporation of CP-AMPARs(Henley & Wilkinson, 2016). Furthermore, we perfused 1-naphthyl acetyl spermine (NASPM; 200 μM), a selective antagonist for CP-AMPARs(Koike *et al*, 1997), but failed to detect any difference in NASPM sensitivities between cKD and control groups regardless of cocaine treatment (Figs. 3B and EV4D). These data indicated that GluN2B ablation facilitated the recruitment of calcium-impermeable AMPARs (CI-AMPARs), but not CP-AMPARs, to D1-MSNs.

**Figure 3.**
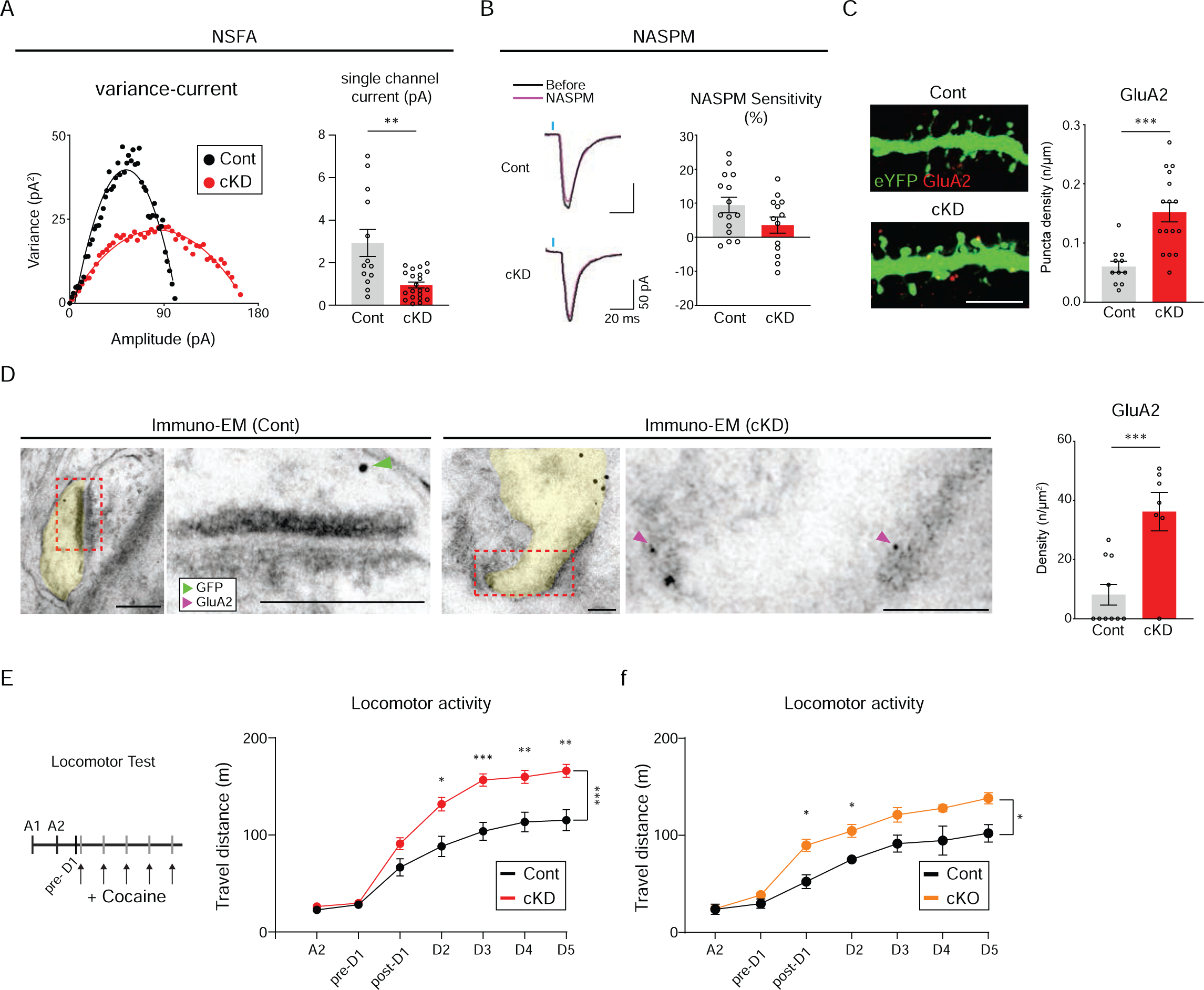
GluN2B ablation facilitated the recruitment of GluA2-AMPARs. (**A)** A variance-current plot from cocaine-treated D1-MSNs (red for cKD; black for control groups, left). Averaged single-channel currents are compared between cocaine-treated mice that previously received either shGluN2B or control virus (n = 21 cells for cKD; n = 13 cells for control groups, right). (**B)** Sample traces of evoked EPSCs before (black) and after (magenta) treatment with NASPM, an antagonist for GluA2-lacking AMPARs in D1-MSNs from cocaine-treated mice that previously received either shGluN2B or control virus (left). The sensitivity to NASPM is compared between cocaine/cKD (n = 13 cells) and cocaine/control (n = 15 cells) groups (right). (**C)** Representative images of GluA2-positive puncta (red) in cocaine-treated D1-MSNs dendrites (green, left). Quantification of GluA2 puncta on D1-MSNs dendrites between cocaine-treated mice that previously received either shGluN2B or control virus (n = 9 cells for cKD; n = 11 cells fo control groups, right). Scale bar = 5 µm. (**D)** Immuno-electron microscopic images for subcellular localization of GluA2 (6 nm gold particles, magenta arrows) and eYFP for labeling of D1-MSNs (18 nm gold particles, green arrows, left). GluA2 particles present within the synaptic areas are compared between cocaine-treated mice that received either shGluN2B or control virus (n = 7 cells for cKD; n = 10 cells for control groups, right). Scale bars = 100 nm. (**E)** An experimental timeline for measurement of cocaine-induced locomotor activity. Locomotor activity was monitored upon cocaine infusion on a daily basis between groups (left). Cocaine-induced locomotion is compared between cKD (red, n =16 mice) and control (black, n = 14 mice) groups (right). (**F)** Cocaine-induced locomotion is compared between cKO (orange, n = 8 mice) and control (black, n = 5 mice) groups. Data are represented as mean ± SEM (error bars); *p < 0.05, **p < 0.01, ***p < 0.001 by two-tailed unpaired t test, Mann-Whitney test, or two-way analysis of variance (ANOVA, with Sidak’s multiple comparisons).

**Figure 4.**
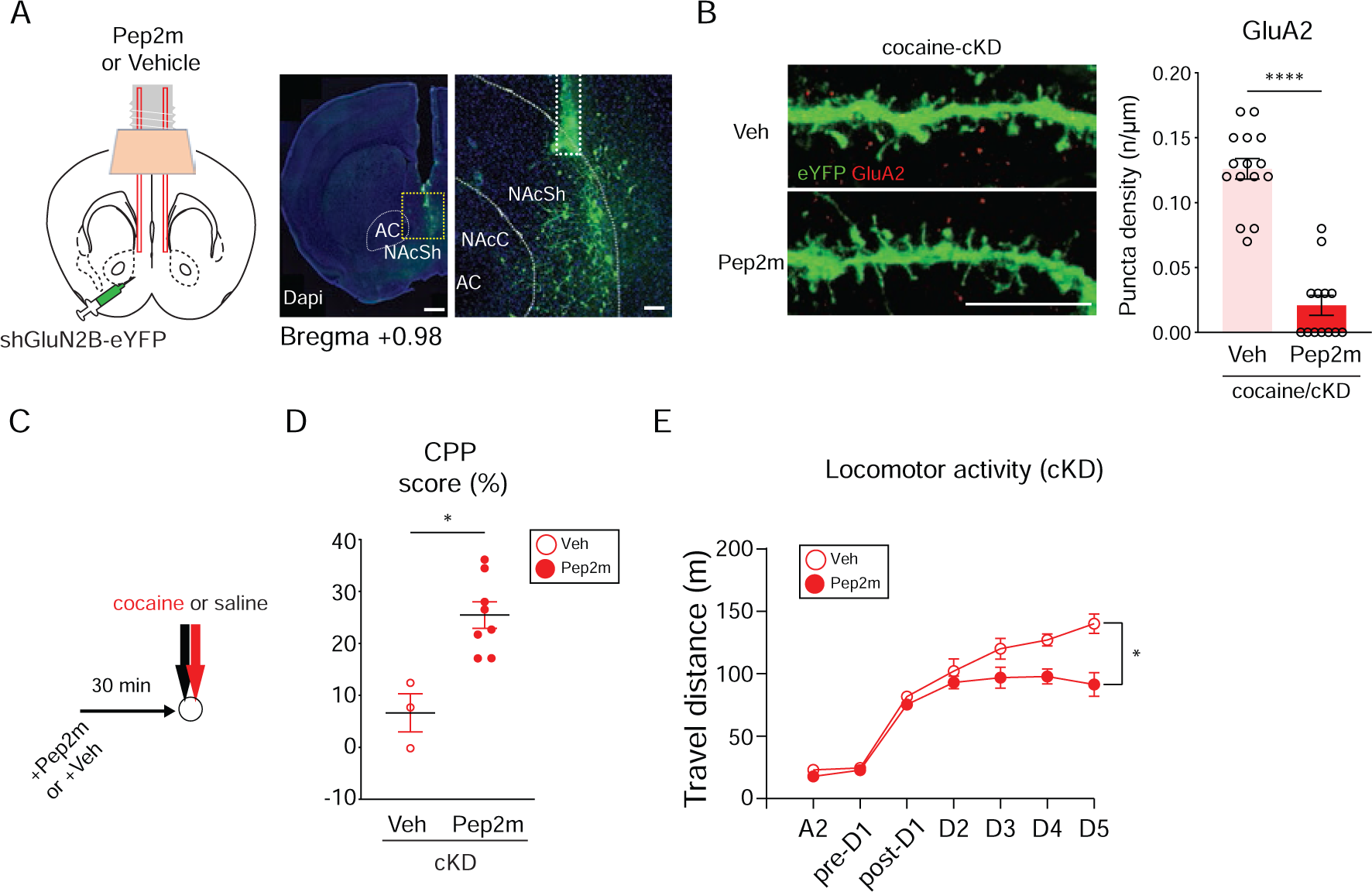
GluN2B depletion induced impairment in addiction memory and elevation of behavioral sensitization attributed to GluA2-AMPAR trafficking**. (A)** A schematic for Pep2m treatment (left). Representative images showing the tips of guide cannulas (dashed line) in the NAcSh (right). Scale bars = 500 μm and 100 μm. (**B)** Representative images for GluA2-positive puncta (red) in dendrites of D1-MSNs (green) from cocaine/cKD mice (left). Scale bar = 10 µm. GluA2 particle density in eYFG-labelled dendrites is compared between GluN2B cKD mice where either Pep2m or vehicle was infused into NAcSh areas (n = 13 cells for Pep2m; n = 15 cells for vehicle groups, right). (**C)** An experimental timeline for behavioral measurement after Pep2m or vehicle infusion. (**D)** Quantified CPP scores in the cocaine chamber after Pep2m or vehicle treatment are compared in mice that previously received shGluN2B virus (n = 8 mice for Pep2m; n = 3 mice for vehicle groups). (**E)** Locomotor activity of GluN2B cKD mice is compared upon daily cocaine infusion after Pep2m or vehicle treatment (n = 5 for Pep2m; n = 3 for vehicle groups). Data are represented as mean ± SEM (error bars); *p < 0.05, ****p < 0.0001 by two-tailed unpaired test, Mann-Whitney test, or two-way analysis of variance (ANOVA, with Sidak’s multiple comparisons).

Given lack of sensitivity changes to NASPM between cKD and control groups as well as no rectification of AMPAR-EPSCs, GluA2-containing CI-AMPARs would be prime candidates for the preferential synaptic recruitment(Henley & Wilkinson, 2016). To verify the incorporation of GluA2-AMPARs, we performed immunohistochemistry (IHC) and immuno-EM using an antibody to GluA2. Our IHC analysis revealed apparent enrichment of GluA2-positive puncta in dendrites of D1-MSNs of cocaine/cKD mice alone but not in other groups (Figs. 3C and EV4E). Immuno-EM also showed an elevated density of GluA2-positive particles within the synaptic areas only in cocaine/cKD group (Figs. 3D and EV4F). Thus, GluA2-AMPARs were likely to be prematurely recruited to D1-MSNs of cKD mice as early as several hours after 5-day exposure to cocaine, which accounted for the precocious maturation of silent synapses of D1-MSNs of cocaine/cKD mice.

Although it was well known that CP-AMPAR incorporation caused behavioral sensitization to cocaine(Terrier *et al*, 2016), whether CI-AMPARs had any similar roles remains unknown. To explore behavioral impacts that the increased incorporation of GluA2-containing CI-AMPARs produced, we compared cocaine-induced locomotor activity of cKD or cKO mice with that of other groups. Indeed, we observed substantial augmentation in locomotion activity to cocaine injection, namely hyper-behavioral sensitization in both cKD or cKO mice while control groups exhibited typical behavioral sensitization (Figs. 3E, F, and EV4G, H). Collectively, the recruitment of GluA2-AMPARs to D1-MSNs appeared to contribute to premature un-silencing of silent synapses and thereby heightened behavioral sensitization to cocaine exposure.

### Cocaine-induced responses restored by blockade of GluA2-AMPAR trafficking

Pep2m interferes with the interaction between the C-terminus of GluA2 and N-ethylmaleimide-sensitive fusion protein, which can disrupt the trafficking of GluA2-AMPARs and reduce their surface expression(Noel *et al*, 1999; Yao *et al*, 2008). We used a myristoylated form of Pep2m, a membrane-permeable peptide inhibitor, to examine the contribution of GluA2-AMPAR trafficking to the physiological and behavioral changes that we have detected. Micro-infusion of Pep2m into the NAcSh (Fig. 4A) effectively reduced surface expression of GluA2 in D1-MSNs of cocaine/cKD group but not in cocaine/control (Figs. 4B and EV5A).

We examined whether the impaired trafficking of GluA2-AMPARs would have any effect on cocaine-induced behaviors that cKD mice exhibited. When cKD mice bilaterally received either Pep2m or vehicle 30 minutes before each cocaine infusion (Fig. 4C), CPP was normalized in cKD mice (Fig. 4D). Furthermore, Pep2m effectively reduced the locomotor activity of cKD mice compared with vehicle-treated cKD mice (Fig. 4E) while it did not affect cocaine-induced locomotion changes of control mice (Fig. EV5B). Taken together, the increased incorporation of GluA2-AMPARs was likely to underlie the abnormal behaviors to cocaine that we have seen in cKD mice.

### Trafficking of GluA2-AMPARs controlled by Rac1 activity

It was demonstrated that synaptic and structural plasticity was dependent on actin dynamics within the synaptic structures(Star *et al*, 2002). For instance, structural and behavioral alterations would be mediated or controlled by Rac1-dependent actin remodeling(Dietz *et al*, 2012; Glebov *et al*, 2015; Wiens *et al*, 2005). Thus, we assessed Rac1 activity in each group of mice through immunolabeling for active GTP-bound Rac1(Dietz *et al*., 2012). Interestingly, D1-MSNs of cocaine/cKD mice had lower Rac1 activity than those of other groups (Figs. 5A and EV5C). We also attempted to optically manipulate the Rac1 activity to examine its roles for cocaine-induced physiological and behavioral changes, using light-activatable Rac1: photo-active Rac1 (PA-Rac1) and photo-insensitive Rac1 (PI-Rac1) as a negative control(Wu *et al*, 2009). After DIO-AAV encoding either PA-Rac1 or PI-Rac1 was micro-infused into D1a-Cre mice, we confirmed the expression and functionality of transduced PA-Rac1 in eYFP-labelled D1-MSNs as revealed by an increase in puncta density of GTP-Rac1 following 473-nm light stimulation (Figs. 5B and EV5D, E).

**Figure 5.**
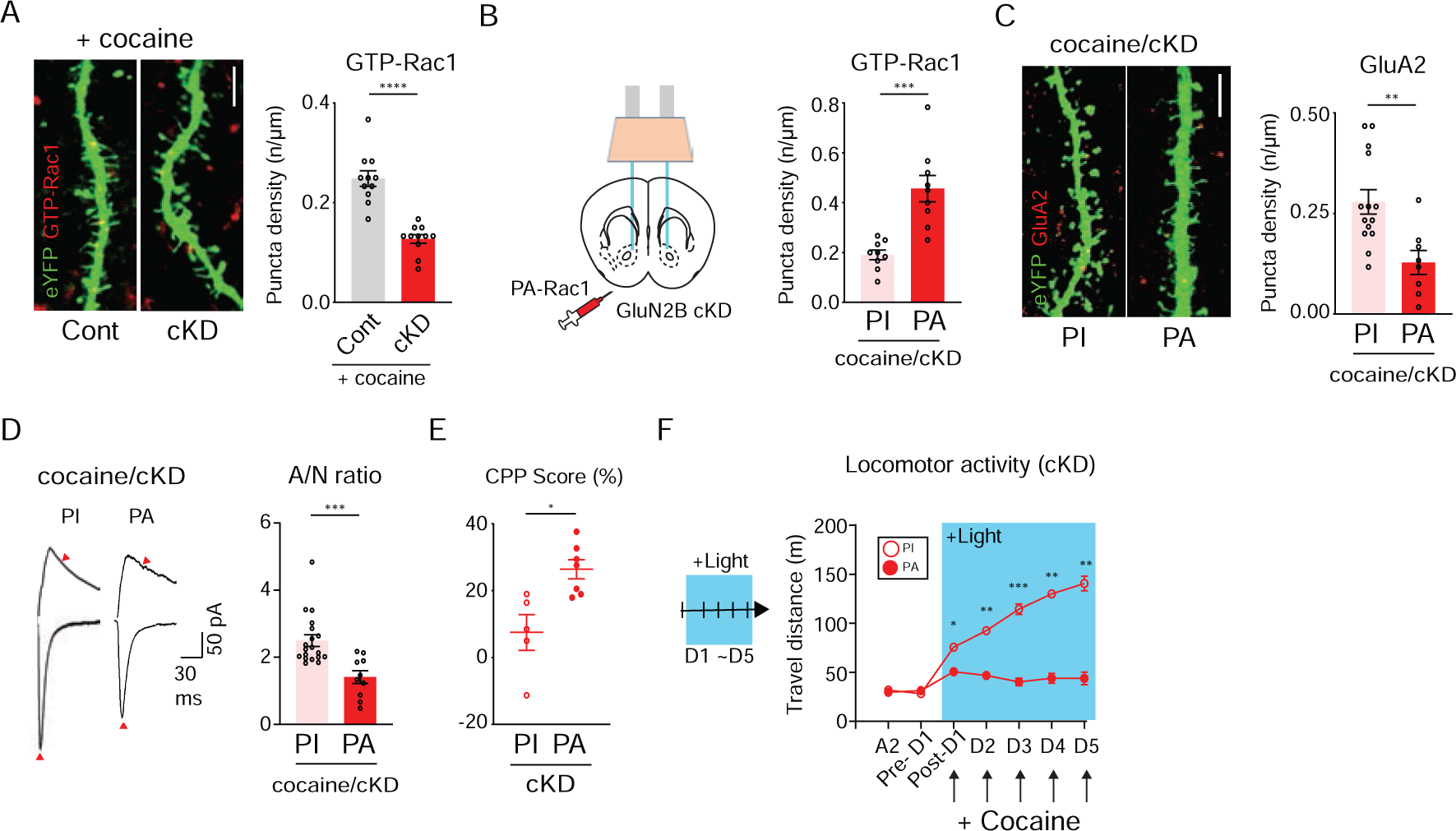
Rac1-controls the trafficking of GluA2-AMPARs. **(A)** Representative confocal images of active Rac1-positive puncta (GTP-Rac1, red) in cocaine-treated D1-MSNs (green, left). Scale bar = 5 μm. Density of GTP-Rac1-positive puncta after cocaine administration is compared between mice that previously received either shGluN2B or control virus (n = 11 cells for cKD; n = 11 cells for control groups, right). (**B)** Experimental schematic for photoillumination-triggered modulation of Rac1 activity (left). GTP-Rac1 puncta density after photoillumination is compared between cocaine/cKD mice that previously received PA-Racl or PI-Rac1 virus (n = 5 cells for PA-Rac1; n = 7 cells for PI-Rac1, right). (**C)** Representative confocal images of active GluA2-positive puncta (red) on the dendrites of cocaine/cKD D1-MSNs (green) expressing PA-or PI-Rac1 (left). Scale bar = 5 μm. Quantified immuno-labeled puncta of GluA2 after photoillumination is compared between cocaine/cKD mice that previously received either PA-Rac1 or PI-Rac1 virus (n = 8 cells for PA-Rac1; n = 14 cells for PI-Rac1 groups, right). (**D)** Sample traces of AMPAR- and NMDAR-EPSCs in cocaine/cKD D1-MSNs after photoillumination of PI or PA-Rac1 (left). A/N ratios are compared after photoillumination between cocaine/cKD mice that received either PA-Rac1 or PI-Rac1 virus (n = 10 cells for PA-Rac1; n = 19 cells for PI-Rac1 groups, right). (**E)** Quantified CPP scores to the cocaine-paired chamber are compared after photoillumination between cKD mice that received either PA-Rac1 or PI-Rac1 virus (n = 9 mice for PA-Rac1; n = 10 mice for PI-Rac1 groups). (**F)** Left: a photoillumination timeline for measurement of locomotor activity. Light was delivered during and after cocaine infusion. Right: cocaine-induced locomotor activity is compared in GluN2B cKD animals that received either PA-Rac1 or PI-Rac1 virus (n = 3 mice for PA-Rac1; n = 4 mice for PI-Rac1 groups). Data are represented as mean ± SEM (error bars); *p < 0.05, **p < 0.01, ***p < 0.001, ****p < 0.0001 by two-tailed unpaired t test, Mann-Whitney, or two-way analysis of variance (ANOVA, with Sidak’s multiple comparisons).

When Rac1 was optically stimulated, the expression of GluA2 decreased in D1-MSNs of cocaine/cKD mice compared with that in PI-Rac1 mice (Figs. 5C and EV5F). This raised the possibility that the physiological and behavioral abnormalities caused by GluN2B ablation could be alleviated by Rac1 activity. Consistent with this notion, A/N ratios obtained from cocaine/cKD mice were normalized by stimulation of Rac1 (Figs. 5D and EV5G), indicating that heightened activity of Rac1 was sufficient to resume synaptic transmission potentially by impeding aberrantly-increased trafficking of GluA2-AMPARs in dendrites of D1-MSNs of cKD mice. We also examined whether activation of Rac1 mitigated the abnormal behaviors that were concurrent with premature incorporation of GluA2-AMPARs. Stimulation of Rac1 restored CPP for the cocaine-paired chamber in PA-expressing cKD mice, whereas we still detected still the impairment of CPP in PI-expressing cKD mice (Fig. 5E). Furthermore, Rac1 activation throughout locomotion tests precluded any behavioral sensitization in cKD and control mice (Figs. 5F and EV5H). Overall, these results underscored Rac1 as a critical regulator that could restrain incorporation of GluA2-AMPARs and abnormal behaviors upon cocaine exposure, which were instigated by GluN2B ablation.

### GluN2B ablation-triggered homeostatic switching of NMDAR subunits

As indicated thus far, GluN2B ablation increased the abundance of GluA2-AMPARs in D1-MSNs of cocaine-treated mice much earlier than previously anticipated by inducing premature un-silencing of silent synapses. However, we noted certain residual silent synapses even after depletion or deletion of GluN2B (Figs. 1E and I), which raised a question which NMDARs contributed to the presence of remaining silent synapses. We compared optically-evoked NMDAR-EPSCs at synapses from the BLA to D1-MSNs at a +50 mV holding potentials in the presence of picrotoxin (PTX) and NBQX (Figs. 6A and EV6A, left). Because the decay kinetics of EPSCs normally depended on the composition of glutamate receptors, we first measured the time taken to decay to half peak amplitude (T_1/2_) rather than the weighted tau(Huang *et al*., 2009). Although we depleted GluN2B that displays slow decay kinetics(Paoletti *et al*, 2013), we still observed slightly longer T_1/2_ in cKD mice than that in control group (Figs. 6A and EV6A, right). The decay kinetics would represent the incorporation of other subunits of NMDARs that had similar or slightly slower decay kinetics than that of GluN2B-NMDARs.

**Figure 6.**
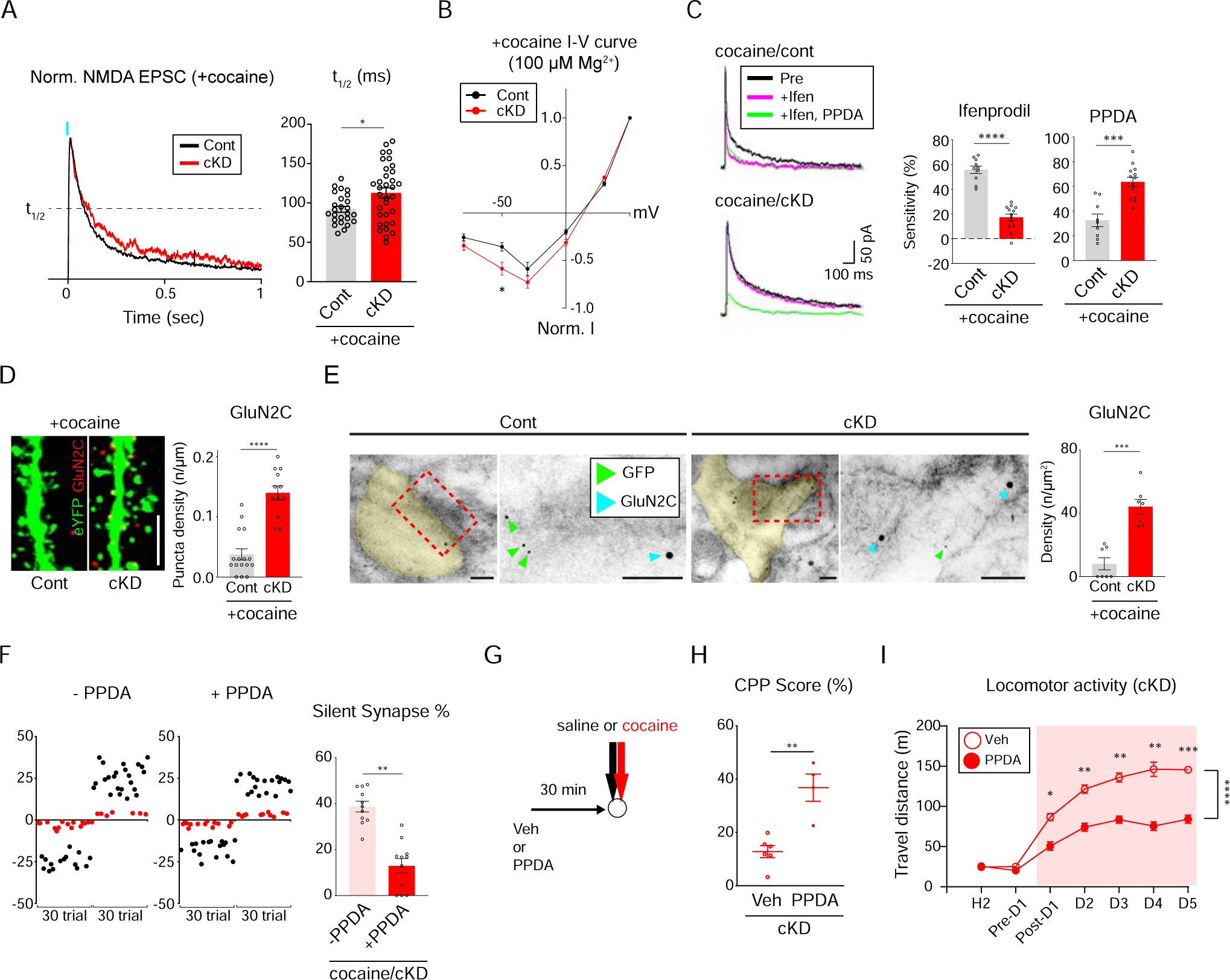
Homeostatic enrichment of GluN2C-NMDARs in the absence of GluN2B. **(A)** Example traces of normalized NMDAR-EPSCs (red traces for cKD; black traces for control, left). Averaged NMDAR-EPSCs from D1-MSNs are compared between cocaine-treated mice that received either GluN2B cKD or control virus (n = 33 cells for cKD; n = 25 cells for control groups, right). (**B)** I-V curves of normalized NMDAR-EPSCs under 100 µM of Mg^2+^ from D1-MSNs in cocaine-treated mice (red, n =18 cells for cKD; black, n = 15 cells for control groups). (**C)** averaged traces of NMDAR-EPSCs from cocaine-treated D1-MSNs with GluN2B antagonist (Ifenprodil, magenta) and additional GluN2C antagonist (PPDA, green) at +50 mV holding potential in the presence of PTX and NBQX (left). Sensitivity of NMDAR-EPSCs to Ifenprodil or PPDA is compared between mice that previously received shGluN2B or control virus (n = 10 cells for Ifenprodil/control; n = 9 cells for PPDA/control; n =13 cells for Ifenprodil/cKD; n = 13 cells for PPDA/cKD, right). (**D)** Representative confocal images of GluN2C-positive puncta (red) in cocaine-treated D1-MSNs (green, left). Scale bar = 5 μm. Quantification of GluN2C puncta after cocaine administration is compared between mice that received either shGluN2B or control virus (n = 12 cells for cKD; n = 16 cells for control groups, right). (**E)** Immuno-electron microscopic images showing subcellular localization of GluN2C (18 nm gold particles, cyan blue arrows) and eYFP for labeling of D1-MSNs (6 nm gold particles, green arrows, left). Scale bar = 100 μm. GluN2C particles present within the synaptic areas are compared between cocaine-treated mice that previously received shGluN2B or control virus (n = 7 cells for both cKD and control groups, right). (**F)** Example plots of evoked EPSCs with minimal optical stimulation (failure trials in red dots; successful trials in black dots) without and with PPDA (left). The proportion of silent synapses is compared before and after PPDA administration in cocaine-treated mice that previously received shGluN2B virus (n = 11 cells, right). (**G)** An experimental timeline of PPDA or vehicle treatment for behavioral assays. (**H)** Quantified CPP scores in the cocaine-paired chamber are compared between PPDA- and vehicle-infused cKD mice (n = 4 mice for PPDA; n = 6 mice for vehicle). (**I)** Locomotor activity was monitored upon cocaine infusion on a daily basis between PPDA- and vehicle-treated cKD mice (n = 4 mice for PPDA; n = 5 mice for vehicle groups). Data are represented as mean ± SEM (error bars); *p < 0.05, **p < 0.01, ***p < 0.001, ****p < 0.0001 by two-tailed unpaired t test, Mann-whitney, Wilcoxon test, or two-way analysis of variance (ANOVA, with Sidak’s multiple comparisons).

Combined with two obligatory GluN1 subunits, other NMDAR subunits, GluN2A, GluN2C, GluN2D, GluN3A, or GluN3B, could be incorporated into silent synapses when GluN2B was absent. To identify subunit configuration of NMDARs in D1-MSNs lacking GluN2B, we perfused 100 µM Mg^2+^ and examined voltage-dependent magnesium blockage of NMDAR-EPSCs. The resultant I-V curves indicated that NMDAR-EPSCs were less sensitive to Mg^2+^ in cKD group than in control group at −50 mV holding potential (Figs. 6B and EV6B). GluN2C/D-containing NMDARs (GluN2C-NMDARs) were less sensitive to Mg^2+^ than GluN2B-NMDARs(Paoletti *et al*., 2013). It was shown that GluN2C-NMDARs had similar decay kinetics to that of GluN2B-NMDARs, GluN2D-NMDARs exhibited much slower decay than NMDARs containing GluN2B or GluN2C while GluN2A-NMDARs showed the fastest kinetics(Paoletti *et al*., 2013; Traynelis *et al*, 2010). Therefore, our physiological data suggested that GluN2C-NMDARs would be highly enriched in NAc D1-MSNs lacking GluN2B and thus were most likely to comprise silent synapses. Consistently, we observed increased expression of GluN2C, but not of GluN2A, from western blots of NAc tissues of cKD mice (Fig. EV6C).

We also performed a sequential antagonism test using specific inhibitors to either GluN2B or GluN2C. The application of Ifenprodil (3 μM) led to significant decreases in NMDAR-EPSC amplitude from cocaine/control mice but not from cocaine/cKD group (Fig. 6c) while Ifenprodil sensitivity did not differ between groups without cocaine treatment (Fig. EV6D). When 0.1 μM of (2S*,3R*)-1-(phenanthrene-2-carbonyl) piperazine-2,3-dicarboxylic acid (PPDA), a well-characterized antagonist for GluN2C/D, was perfused to the same slices, the sensitivity to PPDA was much higher in cKD mice than in control group (Figs. 6C and EV6D). To verify incorporation of GluN2C-NMDARs to D1-MSNs, we performed IHC and immuno-EM using an antibody to GluN2C. Our IHC analysis revealed apparent enrichment of GluN2C-positive puncta in dendrites of D1-MSNs of cKD mice but not in those of control mice regardless of cocaine treatment (Figs. 6D and EV6E). Immuno-EM also revealed a higher density of GluN2C-positive particles in cKD mice compared with that in control mice (Figs. 6E and EV6F). These physiological and anatomical results with the kinetics of observed NMDAR-EPSCs suggested that GluN2B could be replaced by GluN2C in D1-MSNs of mice that repeatedly received cocaine.

It was reasonable to speculate that silent synapses could be installed by GluN2C-NMDARs when GluN2B was absent, leading to aberrant physiological and behavioral consequences. To explore this possibility, we infused PPDA bilaterally into NAcSh areas of cKD mice before each cocaine infusion. PPDA further reduced the proportion of remaining silent synapses of cocaine/cKD mice but did not affect that in other groups (Figs. 6F and EV6G), indicating that GluN2C-NMDARs could contribute to generation of silent synapses in D1-MSNs at least to some extent. Notably, PPDA infusion resumed the typical cocaine-induced behaviors including CPP and behavioral sensitization (Figs. 6G-I), similar to what we had obtained with Pep2m treatment and Rac1 activation. However, PPDA did not affect cocaine-induced locomotor activity in control mice (Fig. EV6H), pointing to negligible contribution of GluN2C without cocaine injection and also PPDA’s specificity to GluN2C-NMDARs.

## Discussion

GluN2B is a requisite component comprising silent synapses, which emerge during the abstinence period after chronic administration of cocaine(Huang *et al*., 2009). However, our knowledge for the functional roles and operational mechanisms of GluN2B-containing silent synapses remains still limited because the *grin2b* gene can be lethal and developmentally effective(Kutsuwada *et al*, 1996). We took advantage of viral-mediated Cre-dependent shRNA and CRISPR-Cas9 strategies to conditionally ablate GluN2B in a region- and cell-type-specific way. Our multidisciplinary investigation by a combination of electrophysiological recordings, cellular and ultrastructural analyses, and behavioral tests, revealed that the ablation of GluN2B reduced the proportion of silent synapses upon cocaine exposure most likely by facilitated recruitment of CI-AMPARs. The resultant AMPAR recruitment-mediated aberrant synaptic potentiation of D1-MSNs resulted in a deficit in CPP, but an elevation of locomotor activity leading to hyper-behavioral sensitization.

Silent synapses in striatal D1-MSNs could play permissive roles in the development of specific responses that appeared after repeated exposure to addictive drugs (Brown *et al*., 2011; Huang *et al*., 2009). In particular, GluN2B has been believed to be a key player in the generation or maintenance of silent synapses since either depletion or antagonism of GluN2B reduced the numbers of cocaine-induced silent synapses(Brown *et al*., 2011; Huang *et al*., 2009) and thereby affected behavioral sensitization and seeking behaviors(Brown *et al*., 2011; Wang *et al*, 2018). However, it was largely unknown how established silent synapses could be sustained after cocaine exposure and which mechanistic roles they exerted for execution of addiction memory and addictive behaviors.

Despite depletion or deletion of GluN2B, we could still detect the residual levels of silent synapses. Our physiological and anatomical analyses indicated that GluN2B was homeostatically replaced by GluN2C which has a similar decay time course of NMDAR-EPSCs to that of GluN2B-NMDARs when GluN2B was ablated in NAc D1-MSNs. Those substituted GluN2C seemed to also yield silent synapses, albeit at a much lower proportion. It was previously shown that the shift of NMDAR subunits would be required for development- and activity-dependent modification of synapses(Barria & Malinow, 2002; Bellone & Nicoll, 2007). In fact, the switching from GluN2B to GluN2C occurred in postnatal cerebellar granule cells(Cull-Candy & Leszkiewicz, 2004; Feldmeyer & Cull-Candy, 1996; Karavanova *et al*, 2007; Ozaki *et al*, 1997). However, it remained unclear how selection and recruitment of GluN2C was made after 5-day exposure to cocaine if it occurred in D1-MSNs lacking GluN2B. Given the state- and activity-dependent changes in the relative abundance and distribution of each GluN2 subunit(Paoletti *et al*., 2013), further investigations for the switching of GluN2 subunits should engender novel insights into mechanisms underlying functional and behavioral consequences that were caused by cocaine-induced silent synapses.

Since Pep2m can interfere with the trafficking of GluA2-AMPARs(Noel *et al*., 1999; Yao *et al*., 2008), the increasement of surface GluA2-AMPARs could be ascribed to aberrantly facilitated trafficking of AMPARs into silent synapses although we could not completely exclude the possibility for the increased production of GluA2. Pep2m-induced normalization of CPP and locomotor activity to cocaine implicated that the enrichment of GluA2-AMPARs would account for abnormal behaviors that mice lacking GluN2B exhibited upon cocaine exposure. These observations prompted us to hypothesize that a deficit in CPP and hyper-behavioral sensitization could result from the heightened excitatory drives that were induced by premature recruitment of GluA2-AMPARs.

Structural alterations such as generation of new synapses and maturation of existing synapses relied on cytoskeletal dynamics (Nakahata & Yasuda, 2018). Trafficking of AMPARs should also involve reorganization of actin cytoskeleton, which allowed for the incorporation of AMPARs into silent synapses(Nakahata & Yasuda, 2018; Star *et al*., 2002). Rac1 is a well-known regulator to modulate actin remodeling in dendrites and spines of neurons(Dietz *et al*., 2012; Wiens *et al*., 2005) and, when suppressed, was able to drive structural plasticity and functional maturation of MSN synapses upon cocaine exposure(Dietz *et al*., 2012; Nestler, 2013). We also detected that puncta numbers of active GTP-bound Rac1 decreased in D1-MSNs of cKD mice compared with those of control mice (Fig. 5A). However, acute stimulation of Rac1 normalized CPP to cocaine but reduced the locomotion activity in cKD mice along with a decreased abundance of surface GluA2 in D1-MSNs, which differed from the previous report indicating that the same stimulation of Rac1 impaired both synaptic maturation and behavioral responses to cocaine(Dietz *et al*., 2012). The apparent discrepancies might be due to different experimental paradigms: D1-MSNs (in this study) vs. general MSNs (Dietz *et al*., 2012)and absence (in this study) vs. presence of GluN2B(Dietz *et al*., 2012). Kalirin 7, an upstream activator of Rac1, could be activated by GluN2B-NMDARs(Kiraly *et al*, 2011)which would control the Rac1-mediated trafficking of AMPARs. Interestingly, KO mice deficient in Kalirin 7 exhibited hyper-behavioral sensitization and impairment of CPP(Kiraly *et al*, 2010), identical to what we observed in GluN2B cKD/cKO mice. Collectively, Rac1 seemed to be involved in the trafficking and specification of recruited AMPARs, although it was unclear how the interplay between Rac1 and AMPARs could be operated.

Surface expression of GluA2 was significantly reduced by perfusion of Pep2m and stimulation of Rac1, both of which interfere with the trafficking of GluA2-AMPARs. These findings supported the notion that the trafficking of CI-AMPARs to silent synapses could be accelerated in the absence of GluN2B. Because maturation/un-silencing of silent synapses was believed to be mediated by incorporation of CP-AMPARs(Lee *et al*., 2013), the facilitated incorporation of GluA2-containing CI-AMPARs was totally unexpected. However, it was possible that other types of AMPARs rather than GluA2-lacking CP-AMPARs could also un-silence cocaine-induced silent synapses and the selection of AMPAR types would be managed by existing NMDAR types residing in those synapses.

It has been previously shown that GluN2B-NMDARs prevented the precocious maturation of developing synapses by blocking premature trafficking of AMPARs(Gray *et al*, 2011). Developmental capability of GluN2B-NMDARs to deter or slow-down the trafficking of AMPARs would also be operative in cocaine-induced silent synapses as if silent synapses that had been a developmental landmark in synapse formation became a critical contributor to addiction memory and incubation(Brown *et al*., 2011; Huang *et al*., 2009; Lee & Dong, 2011). If it were the case, each subunit of NMDARs would have varied efficacies to prevent the recruitment of AMPARs: GluN2B deterred incorporation of AMPARs most effectively, and GluN2A did so less effectively, while GluN2C did so least effectively or even attracted AMPARs. Elucidation of these important issues requires systematic analysis for structural and biophysical features of various types of glutamate receptors.

In this work, we provide for the first-time experimental evidence that GluN2B-NMDARs can specify subunit composition of AMPARs that are subsequently incorporated into silent synapses. Therefore, Glu2B is critically required for the competency of silent synapses to allow for incorporation of CP-AMPARs but inhibit trafficking of CI-AMPARs after cocaine exposure. Since GluN2B can serve as a gateway to forthcoming outcomes of drug-induced silent synapses for addiction memory and the related behaviors, transient and specific inhibitors to GluN2B as well as the suppressive genetochemical interventions would be of great medical and therapeutic interests.

## Methods

### Experimental animals

All procedures followed animal care guidelines approved by POSTECH Institutional Animal Care and Use Committee (Pohang University of Science and Technology, POSTECH-2019-0060). D1a-cre mice were purchased from MMRRC (#034258-UCD) and Rosa26-LSL-Cas9 knock-In (#024857) was from Jackson Laboratory (Bar Harbor, ME). Mice were housed in temperature- and humidity-controlled rooms with a light/dark (12h/12h) cycle with *ad libitum* access to food and water.

### DNA constructs

We employed a suite of shRNA design tools, including BLOCK-It RNAi Designer, siDESIGN Center, and AsiDesigner, for the production of shRNA targeting mouse GluN2B sequence GCTCATCTCTGTGTCATAT. Once designed, shRNAs were confirmed and incorporated into pAAV-Ef1a-DIO-cKD-eYFP-Blank construct, as detailed in earlier studies(Kwon *et al*., 2015; Rubinson *et al*, 2003). For expressing both photo-active (PA) and photo-insensitive (PI) Rac1, we cloned the PA-(Addgene plasmid #22027) and PI-Rac1 (Addgene plasmid #22028) constructs into pAAV-EF1a-double floxed-hChR2(H134R)-eYFP-WPRE (Addgene plasmid #20298)(Wu *et al*., 2009).

To achieve selective expression of *sgGrin2b* sequence in a Cre-dependent fashion, a target sequence, including its PAM site (CGGGACTGTATTCCGCATGCAGG– AGG), was identified using online search tools. This sequence oligomer was then integrated into the pSMART vector (Addgene plasmid #80427) to drive sgRNA transcription under the U6 promoter(Howden *et al*, 2016). The resulting U6-sgGrin2B-scaffold constructs were subsequently cloned into the pAAV-EF1a-DIO-mCherry vector (Addgene plasmid #20299).

### Virus production

Virus production was conducted in accordance with established protocols (Hommel *et al*, 2003). Briefly, HEK-293 cells were co-transfected with helper plasmids (delta-F6), capsid (serotype 5) and pAAV vectors (pAAV-Ef1a-DIO-cKD-eYFP-shGluN2B, pAAV-Ef1a-DIO-cKD-eYFP-Blank, pAAV-Ef1a-DIO-PA-Rac1-mCherry, pAAV-Ef1a-DIO-PI-Rac1-mCherry, pAAV-U6-sgGrin2b-Ef1a-DIO-mCherry, and pAAV-U6-scrambled-Ef1a-DIO-mCherry) at an equal molar ratio using Lipofector-2000 transfection reagents (#AB-LF-2002, AptaBio, Korea). 72 hours after co-transfection, the cells were lysed through freeze-thaw steps and resultant AAV particles were concentrated with Amicon Filter (100K, Millipore, Bedford, MA) to achieve at least 3.0 × 10^12^ gc/ml.

### Animal surgery

Mice were anesthetized using ketamine and xylazine before undergoing surgery using a stereotaxic apparatus (Kopf, Tujunga, CA). 0.3 - 0.5 μl of virus solution was infused using Nanoject II or III (Drummond scientific instrument, Broomall, PA) for 1 minute (23.0 nl per injection, rate of 46 nl/sec), followed by a 10-minute pause for virus diffusion. Coordinates were designated according to the mouse brain atlas (NAcSh: AP +1.35 mm, ML ±0.5 mm, DV - 4.50mm from the bregma; BLA: AP −1.40 mm, ML ±3.35, DV −4.50 mm from the bregma). For micro-infusion of drugs or peptides, bilateral guide cannulas (26-gauge, center-to-center distance: 0.5 mm, Plastics One, Roanoke, VA) were implanted at 0.5 mm above the injection site and fixed on the skull with screws and dental cement. The injection cannula remained capped with a dummy-cannula and dust-cap (Plastics One) after surgery. For *in vivo* optical illumination, optic fibers (NA:0.37; fiber Diameter: 200 μm; ferrule Diameter: φ2.5 mm; fiber Length: 5.0 mm, Newdoon, Zhejiang, China) were implanted at 15° angled with the following coordinates (AP: +1.30 mm, ML ±1.5 mm, DV: −4.53 mm).

### Behavioral analyses

CPP was performed one day after a 5-day cocaine injection as described previously(Tzschentke, 1998). CPP chambers were composed of white or black-lined acryl boxes (20 × 20 cm) and connected with a narrow gray corridor (Fig. EV 1i). Except during acquisition, mice were always placed into the connecting chamber. During acclimation, after a sham intraperitoneal (i.p.) injection, mice explored CPP chambers for 15 minutes. Before the beginning of the test sessions, a preference test was made to determine which chamber would be paired with cocaine or saline (pretest). The preferred place was assigned as saline-paired, while the less preferred place was paired with cocaine injection. If the preference was unclear (time difference < 30 sec), the white box was paired with cocaine and the black with saline. For sucrose CPP, animals were subjected to water deprivation for 3 hours before conditioning sessions. When the conditioning session started, a falcon tube lid containing either a 15 % sucrose solution or plain water was placed. Animals were then given 15 minutes to explore and consume the solution. The CPP score equaled the time difference between the two chambers. Movement paths and durations were analyzed using the SMART 3.0 software (PanLab, Spain), with a mouse’s position determined by its front paw location.

To measure cocaine-induced locomotion, which took place in a 40 x 60 cm white box, mice were given two 10-minute sessions to freely explore and acclimatize. On the next day, after i.p. injection of saline to establish basal activity, mice were treated injected with either saline (0.9%) or cocaine (20mg/kg) at their home cage over 5 consecutive days. On each treatment day, after the injection, mice were placed at the box’s center and their movements were monitored for 10 minutes following a 1-minute acclimation period. The movement trajectories were analyzed using the SMART 2.0 and 3.0 software (PanLab). For photo-illuminated locomotor tests (Figs. 5f and EV 5h), optic fibers connected to a rotary joint hovered above the open field chamber throughout the sessions. Optic fibers were connected to a 473-nm blue laser diode and was continuously turned on during the behavior session with an output of 0.35 mW at the tip of the fibers.

### Electrophysiology

Acute brain slices were prepared for electripysiological recordings within 3 hours after the last injection of cocaine. As described previously(Kwon *et al*., 2015), coronal NAc slices (300 μm thick) were obtained using a vibratome (VT1200s, Leica, Germany) in ice-cold sucrose cutting solution containing (in mM) 20 NaCl, 3.5 KCl, 1.4 NaH_2_PO_4_, 1.3 MgCl_2_, 26 NaHCO_3_, and 11 D-glucose, and 175 sucrose while equilibrated with 95% O_2_ / 5% CO_2_ (pH 7.3 - 7.4). Obtained slices were kept in artificial cerebrospinal fluid (aCSF) containing (in mM) 126 NaCl, 18 NaHCO_3_, 11 D-glucose, 1.6 KCl, 1.2 NaH_2_PO_4_, 1.2 MgCl_2_, and 2.5 CaCl_2_, constantly bubbled with 95% O_2_/ 5% CO_2_ (295∼305 mOsm, pH 7.3 – 7.4) at room temperature (RT) after 30 minutes recovery at 33°C. Slices were placed in recording chambers and continuously perfused (2 ml/min) with aCSF at RT to avoid temperature-dependent increasement of NMDAR-EPSCs^5^, and then whole-cell voltage-clamp configuration was made with a MultiClamp 700B amplifier (Molecular Devices). Recording electrodes (5-7 MΩ; Narshige, Japan) were filled with an internal solution containing (in mM) 117 CsMeSO_4_, 20 HEPES, 0.4 EGTA, 2.8 NaCl, 5 TEA-Cl, 2.5 MgATP, 0.25 Na_3_GTP, and 5 QX-314 (pH 7.2 and 275∼285 mOsm adjusted with CsOH and HEPES, respectively). Recordings were conducted under visual guidance (40 ×, differential interference contrast optics) and expression of eYFP for verification of D1-MNNs was confirmed before recordings. Series resistance of 10-30 MΩ was monitored continuously throughout recordings. Upon changes beyond 15%, any recording data were excluded from further analysis. Synaptic currents were recorded, filtered at 3 kHz, amplified five times, and then digitized at 20 kHz with a Digidata 1322A analog-to-digital converter (Molecular Devices).

We used various chemicals with distinct concentrations as follows, picrotoxin (PTX, 100 μM, Tocris, UK) for blockade of GABAaR-mediated responses, tetrodotoxin (TTX, 1 μM, Tocris) for blockade of sodium channel-mediated action potentials, and 2-amino-5-phosphonovaleric acid (APV, 50 μM, Tocris) for blockade of NMDAR-mediated responses. Ifenprodil hemitartrate (3 μM, #0545, Tocris) and PPDA (0.1 μM, #2530, Tocris) were used to inhibit GluN2B- and GluN2C/D-containing NMDARs, respectively. NBQX (25 μM, #0373, Tocris) was used to inhibit AMPAR-mediated responses. 1-Naplhthylacetyl spermine trihydrochloride (NASPM, 200 μM, #2766, Tocris) was used to selectively inhibit GluA2-lacking CP-AMPARs.

Optical stimulation of axonal fibers of BLA pyramidal neurons to evoke EPSCs was made by illuminating with 0.5-msec blue light pulses from a 470-nm LED source (M470L3, ThorLabs, Newton, NJ). Optically-evoked AMPAR- and NMDAR-EPSCs were measured at −70 mV in the presence of APV and +50 mV in the presence of NBQX, respectively. With 10 - 30 consecutive individual AMPAR-EPSCs, the non-stationary fluctuation analysis (NSFA) was performed with MiniAnalysis (Synaptosoft, Fort Lee, NJ). Decay kinetics of NMDAR-EPSCs were assessed using the time from peak amplitudes of EPSCs to the one-half peak amplitudes ^6,7^.

The mean amplitude of NMDAR-EPSCs was measured by averaging of 10 – 20 consecutives individual EPSCs at various holding potentials (−80, −50, −30, 0, 30 and 50 mV) with the presence of NBQX and PTX. mEPSCs were recorded at −70 mV holding potential in the presence of TTX and PTX for 3 minutes. Detection of mEPSCs was made using a template-based event-detecting module from Clampfit 10.7 software (Molecular Devices).

We used the minimal stimulation method to estimate the proportion of silent synapses as previously described (Brown *et al*., 2011; Huang *et al*., 2009). The frequency of optical stimulation was set at 0.33 Hz. After evoking EPSCs (< 50 pA) at −70 mV, photostimulation intensity was adjusted until both failures and successes could be readily distinguished. The proportion of silent synapses were calculated using the following equation: 1-ln(F_-70_) / ln(F_+50_), in which F_-70_ is the failure rate at −70 mV and F_+50_ is the failure rate at +50 mV.

### Immunohistochemistry (IHC)

Mice were anesthetized by i.p. injection of avertin (250 mg/kg body weight, #T48402, Sigma) and transcardially perfused with phosphate buffered saline (PBS) followed by 4% formaldehyde. Brains was postfixed overnight at 4 °C in a 4 % formaldehyde solution and then embedded in 5 % agarose gel for sectioning (50-µm-thick coronal sections) with a vibratome (VT1000S, Leica).

Sliced sections were blocked with 4 % normal goat serum and 0.45 % Triton X-100 in 0.1 M of phosphate buffer at 4°C for 1 hour and then were incubated with primary antibodies as follows: rabbit anti-GluN2B (AGC-003, Alomone labs., Israel), rabbit anti-GluN2C (AGC-018, Alomone labs.), mouse anti-GluA2 (MAB397, Millipore, MA), rabbit anti-Rac1 (05-389, 1:500, Millipore, MA), or mouse anti-GTP-Rac1 (#26903, NewEast Bioscience, PA) antibodies at 4 °C overnight. Goat anti-rabbit DyLight 650 conjugated IgG (1:500, Bethyl Laboratories, TX), goat anti-rabbit Alexa 568 conjugated IgG (1:500, Invitrogen, CA) or goat anti-mouse DyLight 650 conjugated IgG (1:500, Bethyl Laboratories, TX) were used as secondary antibodies. All tissues were mounted on slide glasses with UltraCruz mounting medium (Santa Cruz Biotech., TX).

### Confocal imaging and laser capture microdissection

We used laser scanning confocal microscopes (LSM 510, Zeiss, Germany and FV3000, Olympus, Japan) for cellular imaging for immuno-labeled puncta co-localization on the dendrites with 40X or 60X objectives. Quantitative analysis of immunoreactive puncta on the dendrites was performed with color threshold (assumed below 80 as a background noise intensity, and the threshold applied green and red signal of pixels as a colocalized) and integrated with MetaMorph 7.7 (Molecular Devices, Sunnyvale, CA).

150 µm-thick coronal brain sections were made with a vibratome and stained with DAPI (4′,6-diamidino-2-phenylindole) to facilitate the alignment of the cell nucleus between confocal and EM images. Brain slices were imaged at low magnification with bright field, GFP and DAPI channels to visualize GFP-labeled neurons with fiducial capillaries and cell nuclei. Following confocal imaging, sections were transferred to a laser capture microdissection system (Zeiss, PALM MicroBeam) for laser etching of the confocal imaged region. The etched sections were trimmed and stored in 2% paraformaldehyde and 2.5% glutaraldehyde in 0.15M cacodylate buffer at 4℃.

### Serial block-face scanning electron microscopy

The brain slices imaged under a confocal microscope were washed in cold 0.15 M sodium cacodylate buffer and placed in cacodylate buffer containing 2% OsO_4_/1.5% potassium ferrocyanide for 1 hr. Tissues were placed in 1% thiocarbohydrazide (TCH, Ted Pella, Redding, CA) in ddH_2_O for 20 minutes and then reacted in 2% aqueous OsO_4_ for 30 min. Thereafter, samples were en bloc stained with 1% uranyl acetate overnight and lead aspartate solution for 30 minutes to enhance membrane contrast as described previously(Walton, 1979). Tissues were dehydrated using an ascending series of ethanol (50%, 70%, 90%, 95% and 100%), and were placed in ice-cold dry acetone. Tissues were then gradually equilibrated with Epon 812 resin (EMS, Hatfield, PA) by placing a mixture of resin and acetone. Samples were embedded in fresh Epon 812 resin containing 7% carbon black (a kind gift from Dr. Nobuhiko Ohno), mounted on aluminum rivets, and cured at 60℃ dry-oven for 2 days for increased specimen conductivity.

A silver paste was applied to the specimen to ground the carbon resin to the aluminum pin. The pin was then coated with 10 nm of gold-palladium in a sputter coater (Q150RS) to further enhance conductivity. Specimens were imaged in a Merlin field emission scanning electron microscope (Carl Zeiss) equipped with 3View2 technology (Gatan) and OnPoint backscattered electron detector (Gatan). Photoshop software was used to overlay the low-magnification EM images atop the confocal images. Dendritic branching patterns of individual GFP-positive neurons were carefully considered to obtain optimal serial EM datasets, including those dendritic profiles. Serial EM images were acquired using an aperture 30 μm, high vacuum, an acceleration voltage of 1.5 kV, an image size of 5,000 by 5,000 pixels, an image resolution (XY plane) of 11 nm, dwell time of 3.5 µs and section thickness of 50 nm. The serial images obtained were converted to 8-bit and were aligned with TrakEM2, an Image J plugin. eYFP-labeled neurons in confocal images were further confirmed with neurons in serial EM image stacks and their dendritic profiles were manually segmented and 3-dimensionally presented.

### Spine classification

Reconstructed dendritic structures were classified with three key parameters (spine length, head width, and neck width). Spine length was manually determined by measuring the maximal length of the entry from the dendrite to the outermost part of the spine head. All dendritic protrusions longer than 0.4 μm were considered for this analysis. For spine width, a line was drawn across the widest part of the dendritic protrusion, including the spine head. In accordance with these parameters, dendritic protrusions were classified into four classes as follows: 1) filopodia had lengths that were larger than the diameters of their necks and heads, similar size of necks to head diameters, and over 3 μm of total extent, 2) thin-shaped spines had over 0.75 μm length of the total extent which were similar to their head and neck diameters. 3) stubby spines had a similar or larger neck diameter than head diameters, and their total extent length was not longer than 0.75 μm. And 4) mushroom spines had diameters (over 0.35 μm) of their heads which were larger than those of their necks. These classes were then categorized into mature (stubby and mushroom) and immature (thin and filopodia) dendritic protrusions for macro-level comparisons. The same parameters were also applied for the analysis of confocal images in Fig. 2 and Fig. EV 3.

### Post-embedding immuno-gold electron microscopy (EM)

We performed transmission electron microscopy (TEM) for immuno-EM imaging with slight adaptation from the established protocol(Masugi-Tokita & Shigemoto, 2007). Transcranial perfusion and preparation of brain slices were identical to those of IHC protocols except for 200 μm thickness. A high-pressure freezing system (HPM 100, Leica) was used to freeze the tissues acutely while preserving membrane and cellular components. Then, sample tissues were kept in acetone and embedded in Lowicryl HM20 resin (Electron microscopy sciences, Hatfield, PA) at −45 °C for 2 days and UV polymerization for 1 day with EM AFS2 (Leica). UV-polymerized blocks containing the NAcSh were sliced by an ultramicrotome (Leica). These slices were then put on the Nickel grids (FCF200-Ni, Electron microscopy sciences, PA). For immunostaining, the same antibodies that were used in IHC experiments (1:50 for GluN2B, GluN2C, and GluA2) as primary antibodies and then paired with anti-GFP (1:50 for mouse monoclonal, sc-9996, Santacruz and rabbit polyclonal, LF-PA0046, AbFrontier, S. Korea) for eYFP detection were utilized. Subsequently, 18 nm gold particle-Donkey anti-rabbit and 6 nm gold particle-Donkey anti-mouse (1:50 for #711-215-152 and #715-195-150, Jackson Immuno Research) antibodies were used for labeling rabbit (for GluN2B, C and GFP) or mouse (GluA2 and GFP) primary antibodies after blocking with 0.2% normal donkey serum in detergent-free PBS at 4°C overnight. For simultaneous staining of GFP, mCherry, and GluN2B, we utilized the same mouse monoclonal anti-GFP antibody and rabbit anti-GluN2B antibody as previously described, with the exception of a rat monoclonal antibody specific for mCherry (1:25, M11217, Invitrogen). Following overnight blocking at 4°C with 0.2% normal goat serum in detergent-free phosphate buffer, we applied 18-nm gold-conjugated goat anti-mouse, 6-nm gold-conjugated goat anti-rat (1:25 for #115-215-146 and #112-195-14, respectively, Jackson ImmunoResearch), and 12-nm gold-conjugated goat anti-rabbit (1:25 for #ab105298, Abcam, UK) secondary antibodies, respectively. After antibody application, we treated 1 - 2% uranyl acetate for 4 minutes and Reynolds solution for 2 minutes to obtain high-contrast images. Images were obtained by transmission electron microscope (JEM-1011, Jeol, Japan) equipped with a CCD camera (ES1000W, 3611 x 2457 pixels, Gatan Inc., CA). To ensure the accuracy of gold particle signals and prevent false positives, consistent observation was maintained without distortion during focus adjustments and X-Y beam alignment.

### Intracranial infusion of drugs or peptides

Mice were bilaterally administered with 0.3 μl of either myristoylated Pep2m (50 μM, #3801, Tocris), PPDA (0.1 μM, #2530, Tocris), or vehicle (aCSF) via an injection cannula connected to a 10 μl Hamilton syringe (Franklin, MA). Micro-infusion was conducted over 3 minutes at a rate of 0.1 μl/min using a micro-infusion pump (Harvard Apparatus, Holliston, MA). Following the injection, mice were allowed a 30 minute rest in their home cages before the start of behavioral tests. The subject mice were transcardially perfused after completion of the last behavioral tests and verified for injection locations. Any mice with incorrectly placed cannula tips were excluded from further analysis.

### Western blotting

Mice were anesthetized with Avertin and transcardially perfused with sucrose cutting solution, and brain slices were made using a vibratome (Leica) as the aforementioned. NAc areas were separated by a steel borer (1.5 mm inner diameter). Tissues were transferred to a homogenization buffer containing 100 ul of RIPA buffer (Sigma) with 10 μl each of the protease inhibitor cocktail (Sigma) and phosphatase inhibitor cocktail (Sigma) and then were sonicated. After sonicating, samples were centrifuged at 13,000 rpm for 15 minutes at 4°C, the supernatant was placed in fresh tubes. Protein concentrations were measured using Bradford protein assay (Bio-Rad Laboratories, Hercules, CA). Samples were prepared with 4x sample buffer (0.25 M Tris base (pH 6.8), 8% SDS, 40% glycerol, 0.002% bromophenol blue, 20% 2-mercaptoethanol), and heat denatured at 37 °C for 30 min. Twenty grams of protein per lane were run in 8% bisacrylamide gel electrophoresis and transferred to PVDF membranes. Membranes were blocked with 5% skim milk in PBST. After blocking, membranes were incubated overnight at 4°C with primary antibodies, mouse anti-GluNR1 (1:1,000 for #05-432, Millipore, MA), rabbit anti-GluNR2A (1:1,000 for #07-632, Millipore, MA), rabbit anti-GluNR2B (1:1,000 for #AGC-003, Alomone lab) or mouse anti-GluN2B (1:500 for #610416, BD Transduction Lab, NJ), rabbit anti-NR2C (1:1,000 for #AGC-018, Alomone lab), rabbit anti-GluA1(1:2,000 for #13185, Cell signaling Technology, MA), mouse anti-GluA2 (1:2,000 for #MAB397, Milipore) and anti-β-actin (1: 10,000 for #A5441, Sigma). Membranes were washed three times for 10 minutes with PBST, and incubated with horseradish peroxidase-conjugated secondary IgGs (1:1,000 for rabbit #170-6515, mouse #170-6516, Bio-Rad, CA or 1:2,000 for #7076s, Cell signaling, MA), for 1hour at RT. Signals from membranes were detected with ECL chemiluminescence (Immoblion Western, Millipore) using a CCD-digital camera system (Amersham Imager 680, Cytiva, MA). Proteins were quantified through ImageJ software, and all the data were normalized to the control proteins.

## Acknowledgements

We appreciate Craig H. Bailey (Columbia University) for critical reading and suggestions. This work was supported by the grants from National Research Foundation of Korea (NRF), NRF 2020R1A6A3A01099508 and NRF 2021R1I1A1A01044359 to H.J.K; NRF 2021R1A6A1A10042944 and POSCO TJ Foundation to S.L; NRF 2020M3H1A1075314 and NRF 2021R1A6A3A01087288 to T.Y; NRF RS-2023-00271562 to H.L; KBRI Basic research program (23-BR-01-03) to K.J.L; NRF 2018R1A3B1052079 and RS-2023-00265883 to J-.H.K.

## Author contributions

H.J.K., S.L., and J-.H.K. conceived the research program and designed experiments. G.H.K. and K.J.L performed SEM-based tomography. H.J.K. and H-.Y.L. obtained and analyzed immuno-EM results. K.S., H-.J.J., H-.Y.L. and J.H.J. contributed to brain surgery, behavioral tests, and data analysis. J.H.P. and T.Y. conducted WB. S.H.L. prepared DNA constructs for conditional depletion/deletion. H.J.K, S.L., and J-.H.K wrote the paper. J-.H.K supervised the entire work.

## The paper explained Problem

Silent synapses that are predominantly comprised of GluN2B are pivotal in the development of addictive behaviors. Despite its importance, specific molecular mechanisms underlying the regulation of silent synapses and their functional contribution remain unclear.

## Results

Our comprehensive studies using viral-mediated Cre-dependent shRNA and CRISPR-Cas9 for cell-type-specific ablation of GluN2B in D1-MSNs reveal that the ablation decreases the proportion of silent synapses, causing premature trafficking of AMPARs, which affects addiction memory formation for cocaine-associated places, but promotes locomotor response. Mechanistically, the recruitment of GluN2B-lacking NMDA receptors elicits synaptic trafficking of calcium-impermeable AMPA receptors (AMPARs), GluA2-AMPARs and the synaptic enrichment of GluA2-AMPARs counteracts memory impairment. Furthermore, the depletion of GluN2B suppresses Rac1 activation, leading to the incorporation of GluA2-AMPARs. The absence of GluN2B also triggers homeostatic regulation of GluN2C-NMDARs, affecting cocaine memory.

## Impact

Our findings represent previously unknown details about GluN2B-containing silent synapses and their roles for molecular and cellular foundations of addiction memory.

## Data Availability

Data and code are available from the corresponding author upon reasonable request.

## Expanded View Figures

**Figure EV1.** Validation of GluN2B depletion. (**A)** A schematic diagram of the used construct and its expression for GluN2B knockdown. (**B)** Western blot analysis of HEK293 cells expressing the designated constructs as well as β-actin as a loading control. (**C)** Western blot analysis of NAc tissues after viral infection. (**D)** A histogram of quantified protein amounts in designated conditions (n = 4 samples each for groups). (**E)** Representative IHC images of GluN2B-positive puncta (red) in D1-MSNs dendrites (green, left). GluN2B puncta density after saline administration is compared between mice that received either shGluN2B or control virus (n = 14 cells for cKD; n = 12 cells for control, right). Scale bar = 5 μm. (**F)** Immuno-EM images showing subcellular localization of GluN2B (18 nm gold particles, blue arrows) and eYFP for labeling of D1-MSNs (6 nm gold particles, green arrows, left). The density of GluN2B-positive particles is compared between saline/cKD and saline/control groups (n = 8 cells for cKD; n = 10 cells for control, right). Scale bars = 100 nm. (**G)** Verified location of microinjection sites in the NAc. Brain images (adapted from Paxinos and Watson, 2001)(Paxinos *et al*, 2001) corresponding to the NAc are overlapped, and injection sites are represented by black dots. Injector sites are located in brain slices at indicated distances (mm) from the bregma. **(H)** Example plots of optical responses with minimal optical stimulation in the designated groups (failure trials in red dots; successful trials in black dots, left). The proportion of silent synapses is compared between saline-treated mice that previously received either shGluN2B or control virus (n = 28 cells for cKD; n =19 cells for control, right). (**I)** An actual behavioral setting for CPP (left). CPP was tested before cocaine exposure (n = 10 mice for cKD; n = 17 mice for control, right). Data are represented as mean ± SEM (standard error of the mean); *p < 0.05, ****p < 0.0001 by two-tailed Mann-Whitney or unpaired t test.

**Figure EV2.** Validation of GluN2B deletion. (**A)** Western blot analysis of NAc tissues of mice that received either sgGrin2b or control virus (n = 3 samples each). (**B)** Immuno-EM images showing subcellular localization of GluN2B (18 nm gold particles, blue arrows) and eYFP for labeling of D1-MSNs (6 nm gold particles, green arrows, left). The density of GluN2B-positive particles is compared between saline/cKD and saline/control groups (n = 8 cells for cKD; n = 10 cells for control, right). Scale bars = 100 nm. (**C)** An experimental configuration for sucrose CPP (left). CPP scores to the sucrose-paired chamber are compared between cKO and control mice (n = 6 mice each for control and cKO groups, right). Data are represented as mean ± SEM (standard error of the mean); ****p < 0.0001 by two-tailed unpaired t test or Mann-Whitney test.

**Figure EV3.** Structural and functional features of D1-MSNs in the absence of GluN2B. (**A)** Spine density of D1-MSNs is compared between cKD and control mice that received saline (n = 29 cells for cKD; n = 21 cells for control, left). Spine density of D1-MSNs is compared between cKD and control mice that repeatedly received cocaine (n = 32 cells for cKD; n = 16 cells for control, right). (**B)** Representative illustrations of reconstructed dendrites and protrusions in saline-treated D1-MSNs (top). The proportion of morphologically-categorized spine classes between saline-treated mice that previously received either shGluN2B or control (n =66 spines from 3 mice for cKD; n = 113 spines from 3 mice for control, bottom). (**C)** Structural parameters (spine volume, SV and density of perforated PSD) are compared between cocaine-treated mice that previously received either shGluN2B or control virus (spine volume and SV: n = 95 units for cKD; n = 85 units for control, density of perforated PSD: n =4 units each). (**D)** Structural parameters after saline exposure are compared between mice that previously received either shGluN2B or control virus (head width, spine volume, SV, and PSD area: n = 66 units for cKD; n = 130 units for control; density perforated PSD and branched spines: n = 5 units for cKD; n = 6 units for control). (**E)** Sample traces of mEPSCs from D1-MSNs of saline-treated animals that previously received either shGluN2B or control virus. (**F)** A cumulative probability plot of mEPSC inter-event intervals (IEIs) and an inserted summary histogram of mEPSC frequency after saline exposure (n = 22 cells for cKD; n = 30 cells for control). (**G)** A cumulative probability plot of amplitude after saline exposure and an inserted summary histogram of mEPSC amplitude (n = 22 cells for cKD; n = 30 cells for control). (**H)** Sample traces of optically-evoked EPSCs with stimulation of 50-ms interval in D1-MSNs of saline-treated mice that previously received either shGluN2B or control virus (left). Measurement of PPRs with stimulation of 50-, 100-, and 150-ms intervals in D1-MSNs of saline-treated mice that previously received either shGluN2B or control virus (n= 14 cells for cKD; n = 10 cells for control, right). **(I)** Sample traces of optically-evoked AMPAR-(at −70 mV) and NMDAR-EPSCs (+50 mV) in D1-MSNs of saline-treated mice that previously received either shGluN2B or control virus (left). A/N ratios are compared between saline-treated mice that received either shGluN2B or control virus (n = 17 cells for cKD; n =18 for control, right). Data are represented as mean ± SEM (standard error of the mean).

**Figure EV4:** Comparative analysis of AMPAR features. (**A)** AMPAR I/V curves in cocaine-treated D1-MSNs (left). The rectification indices are compared between cocaine-treated mice that previously received either shGluN2B or control virus (n = 12 for cocaine/cKD; n =11 cells for cocaine/control, right). (**B)** AMPAR I/V curves in saline-treated D1-MSNs (left). The rectification indices are compared between saline-treated mice that previously received either shGluN2B or control virus (n = 12 for saline/cKD; n =13 cells for saline/control, right). (**C)** Representative variance-current plots for saline-treated D1-MSNs (left). Averaged single-channel currents are compared between saline-treated mice that previously received either shGluN2B or control virus (n = 17 cells for saline/cKD; n = 18 cells for saline/control, right). (**D)** Sample traces of EPSCs before (black) and after (magenta) NASPM treatment in saline-treated D1-MSNs (left). Sensitivity to NASPM is compared between saline-treated mice that previously received either shGluN2B or control virus (n = 15 cells for saline/cKD; n = 14 cells for saline/control, right). (**E)** Representative IHC images for GluA2-positive puncta (red) on eYFP-labelled dendrites of saline-treated D1-MSNs (green, left). Density of GluA2-positive puncta is compared between cKD and control groups (n = 23 cells for saline/cKD; n = 9 cells for saline/control, right). Scale bar = 5 µm. (**F)** Representative immuno-EM images showing subcellular localization of GluA2 (6 nm gold particles, magenta arrows) and eYFP for labeling D1-MSNs (18 nm gold particles, green arrows, left). Density of GluA2-positive particles is compared between saline-treated mice that previously received either shGluN2B or control virus (n = 8 cells for saline/cKD; n = 6 cells for saline/control, right). Scale bar = 100 nm. (**G)** a schematic timeline for locomotion measurement (left). saline-induced locomotor activity is compared between mice that received either shGluN2B or control virus (n = 14 for cKD; n = 12 mice for control, right). (**H)** Saline-induced locomotor activity is compared between Cas9-eGFP mice that received either *sgGrin2b* or the control virus (n = 5 mice each). Data are represented as mean ± standard error of the mean (standard error of the mean); *p < 0.05, **p < 0.01 by unpaired t test, Mann-Whitney test, two-way analysis of variance (ANOVA, with Sidak’s multiple comparisons).

**Figure EV5.** Rac1 mediates the regulation of GluA2 enrichment and cocaine-associated behaviors. (**A)** Representative IHC images for GluA2-positive puncta (red) in dendrites of cocaine/control D1-MSNs (green, left). Density of GluA2-positive puncta is compared between cocaine/control mice that were infused with either Pep2m or vehicle (n = 15 for Pep2m; n = 11 cells for vehicle, right). Scale bar = 5 µm. (**B)** Cocaine-induced locomotor activity is compared between control mice that were infused with either vehicle or Pep2m (n = 5 mice for Pep2m; n = 3 mice for vehicle). (**C)** Representative IHC images of GTP-Rac1-positive puncta (red) in dendrites of saline-treated D1-MSNs (green, left). Density of GTP-Rac1-positive puncta is compared between saline-treated mice that previously received either shGluN2B or control virus (n = 16 cells for saline/cKD; n = 12 cells for saline/control, right). Scale bar = 5 μm. (**D)** Representative fluorescence images for PA-Rac1 or PI-Rac1 expression (cyan) in dendrites of D1-MSNs (green, left). Scale bar = 50 μm. (**E)** The density of GTP-Rac1-positive puncta after photoillumination is compared between cocaine-treated control mice that received either PA-Rac1 or PI-Rac1 (n = 11 cells for PA-Rac1; n = 17 cells for PI-Rac1, right). Scale bar = 50 μm. (**F)** Representative IHC images of GluA2-positive puncta (red) on dendrites of cocaine/control D1-MSNs (green) expressing either PA-or PI-Rac1 (left). density of GluA2-positive puncta is compared between cocaine/control mice that received either PA-Rac1 or PI-Rac1 virus (n = 8 cells for PA-Rac1; n = 7 cells for PI-Rac1, right). Scale bar = 5 μm. (**G)** Example plots of EPSCs evoked with minimal optical stimulation in designated groups. The measurement points for AMPAR-(bottom) or NMDAR-EPSCs (top) are marked with red arrowheads (left). A/N ratios after photoillumination are compared between cocaine/control mice that received either PA-Rac1 (n = 19 cells) or PI-Rac1 virus (n = 10 cells, right). (**H)** Cocaine-induced locomotor activity with photoillumination is compared between control mice that receive either PA-Rac1 or PI-Rac1 virus (n = 3 mice each). Data are represented as mean ± SEM (standard error of the mean); *p < 0.05 by two-tailed Mann-Whitney test.

**Figure. EV6.** GluN2C contributes to silent synapses in the absence of GluN2B. **(A)** Example traces of normalized NMDAR-EPSCs from saline-treated control (black) and cKD (red) D1-MSNs (left). Times taken to the half peak of NMDAR-EPSCs are compared between saline-treated mice that receive either shGluN2B or control virus (n = 25 cells for cKD; n = 28 cells for control, right). (**B)** I-V curves of normalized NMDAR-EPSCs under 100 µM of Mg^2+^ in saline-treated D1-MSNs (n = 14 cells each). (**C)** Western blot analysis for GluN2A and GluN2C expression with biopsy samples of the NAc (left). Western blot data are quantified and compared between mice that receive either shGluN2B or control virus (n = 4 samples each, right). (**D)** Example traces of NMDAR-EPSCs in saline-treated D1-MSNs with sequential application of a GluN2B antagonist (Ifenprodil, magenta) and a GluN2C/D antagonist (PPDA, green) at +50 mV holding potential (left). Sensitivity to Ifenprodil or PPDA is compared between saline-treated mice that received shGluN2B (n =10 cells for Ifenprodil; n = 11 cells for PPDA) or control virus (n = 8 cells for Ifenprodil; n = 10 cells for PPDA, right). (**E)** Representative IHC images of GluN2C-positive puncta (red) on dendrites of saline-treated D1-MSNs (green, left). Density of GluN2C-positive puncta is compared between saline/cKD and saline/control D1-MSNs (n = 11 cells for saline/cKD; n = 21 for saline/control, right). (**F)** Immuno-EM images showing subcellular localization of GluN2C (18 nm gold particles, blue arrows) and eYFP for labeling of D1-MSNs (6 nm gold particles, green arrows, left). Density of GluN2C-positive particles is compared between saline-treated mice that previously received either shGluN2B or control virus (n = 8 cells for cKD; n = 7 cells for control, right). Scale bar = 100 μm. (**G)** Example plots of evoked EPSCs with minimal optical stimulation (failure trials in red dots; successful trials in black dots) without and with PPDA (left). The proportion of silent synapses is compared before and after PPDA administration in cocaine-treated mice that previously received control virus (n = 11 cells each, right). (**H)** Cocaine-induced locomotor activity is compared between control mice that were infused with either PPDA or vehicle (n = 3 mice for PPDA; n = 5 mice for vehicle). Data are represented as mean ± SEM (standard error of the mean); *p < 0.05, **p < 0.01, ***p < 0.001, ****p < 0.0001 by two-tailed unpaired test or Mann-Whitney test.

